# Activation loop dynamics are controlled by conformation-selective inhibitors of ERK2

**DOI:** 10.1101/639567

**Authors:** Laurel M. Pegram, Jennifer C. Liddle, Yao Xiao, Maria Hoh, Johannes Rudolph, Dylan B. Iverson, Guy P. Vigers, Darin Smith, Hailong Zhang, Weiru Wang, John G. Moffat, Natalie G. Ahn

**Affiliations:** Department of Biochemistry, University of Colorado, Boulder, CO 80305; Array BioPharma Inc., 3200 Walnut St., Boulder, CO, 80301; Genentech, Inc., 1 DNA Way, South San Francisco, CA, 94080.

**Keywords:** ERK2/MAPK1, DUSP6, kinase, dynamics, conformational selection, NMR, X-ray crystallography, hydrogen exchange mass spectrometry, Vertex-11e, SCH772984, GDC-0994

## Abstract

Modulating the dynamics of protein kinases expands the inhibitory mechanisms for small molecules. NMR measurements of the MAP kinase, ERK2, have shown that activation by dual-phosphorylation induces global motions involving exchange between two states, “L” and “R”. We show that ERK inhibitors Vertex-11e and SCH772984 exploit the small energetic difference between L and R to shift the equilibrium in opposing directions, while inhibitor GDC-0994 and ATP analogue AMP-PNP retain L⇌R exchange. An X-ray structure of active 2P-ERK2 complexed with AMP-PNP reveals a shift in the Gly-rich loop along with domain closure to position the nucleotide in a more catalytically productive conformation relative to inactive 0P-ERK2:ATP. X-ray structures of 2P-ERK2 complexed with Vertex-11e or GDC-0994 recapitulate this closure, which is blocked in a complex with a SCH772984 analogue. Thus, the L→R shift in 2P-ERK2 is associated with movements needed to form a competent active site. Solution measurements by hydrogen-exchange mass spectrometry (HX-MS) reveal distinct binding modes for Vertex-11e, GDC-0994 and AMP-PNP to active *vs* inactive ERK2, where the extent of HX protection matches their degree of R-state formation. In addition, Vertex-11e and SCH772984 show opposite effects on HX near the activation loop, suggesting that L⇌R exchange involves coupling between the activation loop and the active site. Consequently, these inhibitors differentially affect MAP kinase phosphatase activity towards 2P-ERK2. We conclude that global motions in ERK2 promote productive nucleotide binding, and couple with the activation loop to allow control of dephosphorylation by conformation-selective inhibitors.

**SIGNIFICANCE STATEMENT:** Protein kinases in the RAF/MKK/ERK signaling pathway are dysregulated in cancer and are important targets for inhibitor development. Catalytic activation of the MAP kinase, ERK2, induces global motions involving exchange between two conformational states. Using nuclear magnetic resonance (NMR) and hydrogen-exchange mass spectrometry, we show that inhibitors exploit these motions to trap ERK2 in distinct states. Our findings reveal motions of the activation loop coupled to the active site. Inhibitor binding can control these activation loop dynamics to alter its rate of dephosphorylation by MAP kinase phosphatase.

## INTRODUCTION

Extracellular-regulated kinases (ERK1/2) are key enzymes in the MAP kinase pathway, downstream of A/B/C-RAF and MAP kinase kinase-1/2 (MKK1/2, *aka* MEK1/2). ERKs are tightly regulated, with full activation following dual phosphorylation by MKK1/2, and inactivation following dephosphorylation by MAP kinase phosphatases (e.g. MKP3/DUSP6). The prevalence of oncogenic BRAF mutations in melanoma and other cancers has led to the development of inhibitors targeting BRAF-V600E/K and MKK1/2 (1). While these inhibitors have achieved dramatic success in clinic for metastatic melanoma, many patients develop resistance, often due to reactivation of ERK signaling (1). Because of this, ERKs are potential targets for therapeutics, with high-affinity inhibitors in early-stage clinical trials (2,3). Therefore, understanding regulatory mechanisms for ERKs and how they impact inhibitor binding is a timely and important goal.

Crystal structures of ERK2 in its inactive, unphosphorylated (0P) and active, phosphorylated (2P) forms show a major conformational rearrangement of the activation loop following phosphorylation at regulatory sites, T183 and Y185 (numbered as in rat ERK2, Fig. S1) (4,5). Other conformational changes include a small rotation and closure between N- and C-terminal domains and exposure of a docking motif binding site below the activation loop (4–6). In other kinases, phosphorylation and remodeling of the activation loop cause significant structural changes that position active site residues for catalysis (7). Curiously, the positions of these catalytic residues differ little between 0P- and 2P-ERK2 and resemble those seen in active conformations of most protein kinases (8). This suggests that additional factors are needed to explain why 0P-ERK2 is inactive until phosphorylated.

Solution measurements of ERK2 have revealed changes in protein dynamics following kinase activation. Studies using hydrogen-deuterium exchange mass spectrometry (HX-MS) showed that phosphorylation of ERK2 alters rates of hydrogen exchange in localized regions, which can be ascribed to changes in conformational mobility (9). NMR (^13^C,^1^H) multiple quantum Carr-Purcell-Meiboom-Gill relaxation dispersion measurements on ERK2 selectively labeled with [methyl-^1^H,^13^C]Ile, Leu, and Val revealed global motions throughout the kinase core upon activation by phosphorylation (10,11). Global exchange could be modeled by an equilibrium between two conformational states, named “L” (locked) and “R” (released), interconverting on a millisecond timescale. Thus, allosteric regulation of ERK2 involves constraint of the unphosphorylated kinase in the L state, and formation of the R state upon phosphorylation and associated activation. The conformational changes accompanying L⇌R interconversion and how they compare to those of other kinases have not been elucidated.

Clues to defining R and L have been provided by small molecules with differential binding properties to 0P- and 2P-ERK2. For example, the pyrimidylpyrrole inhibitor, Vertex-11e (12), has 7-fold higher affinity for 2P-ERK2 over 0P-ERK2 (Table S1) and shifts the L⇌R equilibrium in 2P-ERK2 completely to the R state (13). In a second example, the non-hydrolyzable nucleotide, AMP-PNP, leads to greater protection of the catalytic site from solvent D_2_O in 2P-ERK2 than in 0P-ERK2, as measured by HX-MS (14,15) Such findings suggest that the L⇌R equilibrium in 2P-ERK2 allows conformational selection, a property characteristic of Type II kinase inhibitors (16). Type II inhibitors recognize the conserved DFG motif, which coordinates Mg^2+^-ATP. They selectively bind to an inactive “DFG-out” conformation, which is in exchange with the active “DFG-in” conformation. Such conformation selectivity may shift exchanging populations of a kinase into one major state. However, current X-ray structures of ERK2 show no evidence for a DFG-in/DFG-out switch (4,5). The ways in which conformation-selective inhibitors affect 2P-ERK2 structure and dynamics, their relationship to DFG-in/DFG-out states, and their impact on regulation, are currently unresolved.

The goal of this study is to characterize the L⇌R equilibrium in 2P-ERK2 and its impact on kinase regulation. Solution (NMR, HX-MS) and structural (X-ray) measurements are integrated to examine interactions between ERK2 and inhibitors (Vertex-11e, GDC-0994, and SCH772984/SCH-CPD336; Fig. S2) or AMP-PNP. We demonstrate that both Vertex-11e and SCH772984 show properties of conformational selection when bound to 2P-ERK2, shifting the L⇌R equilibrium in opposite directions to R and L, respectively, while GDC-0994 and AMP-PNP binding allow equilibrium exchange, in a manner favoring R. X-ray structures of ERK2 complexed with nucleotide reveal that the L→R transition is associated with a shift from nonproductive to productive nucleotide binding mode. In contrast, structures of 2P- and 0P-ERK2 complexed with the high-affinity inhibitors are identical within the active site. No evidence for DFG-out conformers are observed under any condition. However, HX-MS measurements reveal differential interactions of Vertex-11e and GDC-0994 within the active sites of 2P- *vs* 0P-ERK2 and correlations between degree of HX protection and formation of the R state. Furthermore, HX-MS suggests that the conformation-selective properties of Vertex-11e and SCH772984 affect motions of the activation loop. In accordance, phosphatase-catalyzed dephosphorylation of ERK2 is inhibited by Vertex-11e and increased by SCH772984. The results reveal that motions within the catalytic site promote nucleotide binding for catalytic competency, and are coupled to activation loop motions which control recognition and feedback by downstream enzymes.

## RESULTS

ERK2 selectively labeled with [methyl-^1^H,^13^C] Ile, Leu, and Val (ILV) was examined by NMR before and after activation by *in vitro* phosphorylation. Previous Carr-Purcell-Meiboom-Gill (CPMG) NMR relaxation dispersion experiments showed that 2P-ERK2 exists in equilibrium between R and L states with a population ratio of 80:20, reflecting a difference in free energy of ~0.8 kcal/mol at 25 °C (11). The relative populations of these states can be estimated by heteronuclear multiple-quantum coherence (HMQC) spectra of slow exchanging side chain methyls where chemical shift differences between R and L (in Hz) are larger than the global chemical exchange rate constant (i.e., Δω≫ k_ex_). This is illustrated in Fig. 1A, where methyls corresponding to I72, L242, and L288 each appear as two overlapping peaks in apo 2P-ERK2, consistent with 80:20 populations of R:L states. In contrast, methyls in 0P-ERK2 appear as single peaks, corresponding to only the L state.

**Fig. 1:**
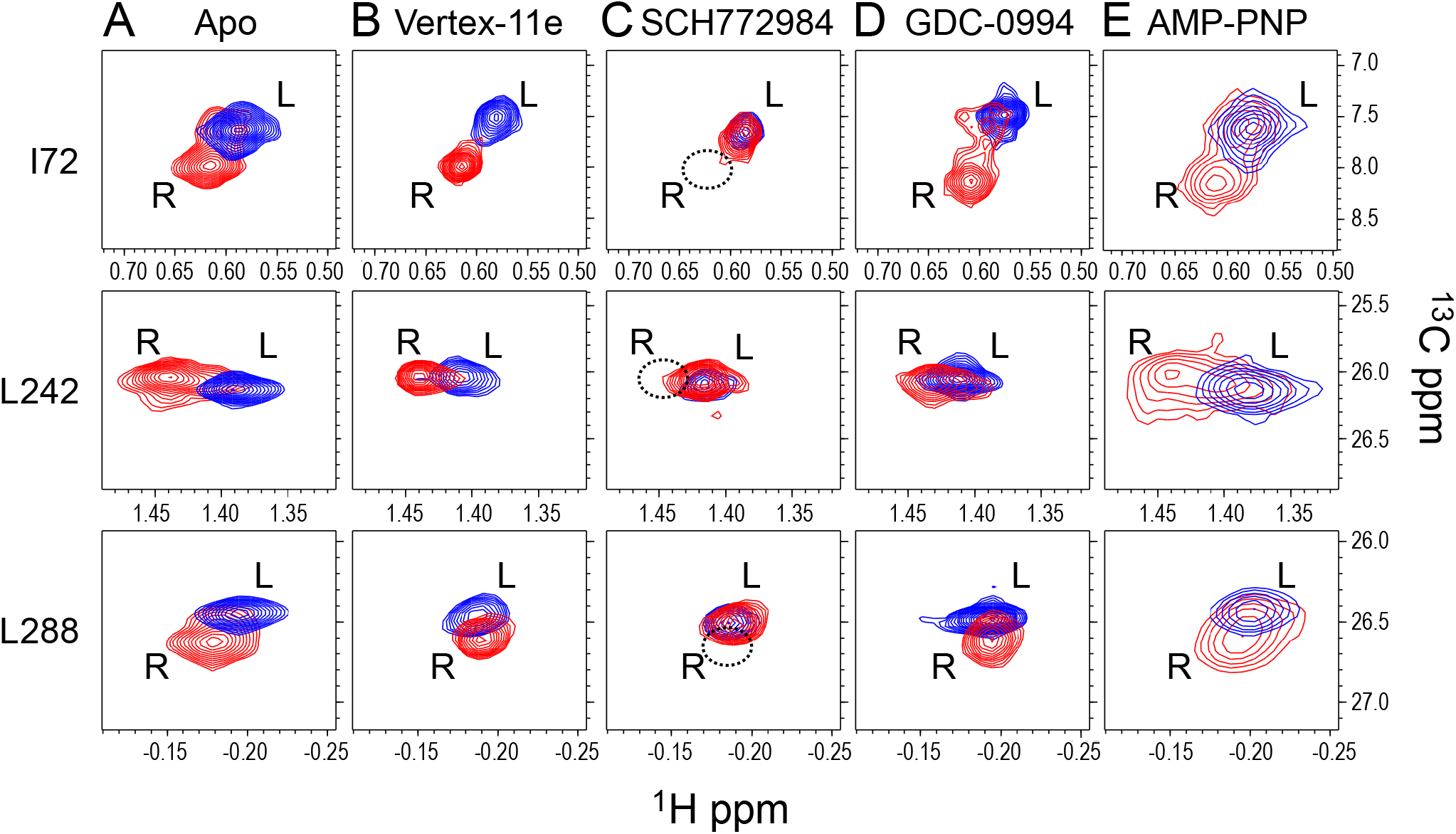
Conformation selection by high affinity ERK inhibitors. Two-dimensional (^13^C−^1^H) methyl HMQC spectra show examples of (^13^C,^1^H)-labeled ILV methyl peaks in 0P-ERK2 (blue) and 2P-ERK2 (red) at 25 °C. **(A)** In ERK2 apoenzymes, each methyl appears as a single peak in 0P-ERK2 corresponding to L, and two overlapping peaks in 2P-ERK2 corresponding to ~20% L and ~80% R states in slow conformational exchange. **(B)** Vertex-11e complexed with 2P-ERK2 causes a population shift to the R state, while 0P-ERK2 remains in the L state. **(C)** SCH772984 complexed with 2P-ERK2 shifts the population to the L state, while 0P-ERK2 remains in the L state. **(D)** GDC-0994 complexed with 0P- or 2P-ERK2 yields R and L state ratios similar to apoenzymes. **(E)** AMP-PNP complexed with 0P- or 2P-ERK2 yields R and L state ratios similar to apoenzymes.

We used this method to determine R:L population ratios for ERK2 complexed with three nanomolar affinity inhibitors, Vertex-11e, SCH772984, and GDC-0994 (Fig. S2, Table S1) (12,17,18). When complexed with Vertex-11e, the [^13^C,^1^H] methyls shifted completely to the R state in 2P-ERK2 (Fig. 1B) as described (13). In contrast, SCH772984 complexed with 2P-ERK2 resulted in a complete shift to the L state (Fig. 1C). Binding of GDC-0994 as well as the ATP analogue, AMP-PNP, to 2P-ERK2 resulted in R:L ratios similar to apoenzyme (Fig. 1D,E). None of the ligands altered the dominant L state in 0P-ERK2. The results reveal that various ligands exploit the L⇌R equilibrium in 2P-ERK2 in different ways. In particular, Vertex-11e and SCH772984 confer conformational selection for R and L states, respectively, while GDC-0994 and AMP-PNP maintain the equilibrium exchange observed in apo 2P-ERK2.

### Distinct binding modes for ERK2 complexed with nucleotide

In order to examine possible conformational differences between R and L states, we determined an X-ray structure of 2P-ERK2 co-crystallized with Mg^2+^-AMP-PNP (Fig. 2A, Table S2, PDBID:6OPG). Two Mg^2+^ ions occupy protein kinase metal positions Me1 and Me2 (19), and bridge the nucleotide P_γ_-oxygen atoms with N152 and D165, placing P_γ_ in proximity to the catalytic base, D147. This differs significantly from a published structure of 0P-ERK2 complexed with ATP (Fig. 2B) (20). The structures show striking differences in the positioning of the ribose ring, where nucleotide dihedral angles between ribose and adenine (O4’-C1’-N9-C8 and C2’-C1’-N9-C4) each differ by 50 degrees. In the inactive complex, P_γ_ coordinates with only one bound Mg^2+^ (Me2) and overlaps the position of P_β_ in 2P-ERK2:AMP-PNP, placing the P_γ_-oxygens more than 6 Å from D147. In contrast, the P_γ_-oxygens are 3.9 Å from D147 in the active 2P-ERK2:AMP-PNP complex. Thus, the active ERK2 complex represents a productive nucleotide binding mode. Similar differences in ribose orientation can be seen in reported structures of 2P-ERK2 complexed with AMP-PCP (PDBID:5V60; 21) and 0P-ERK2 complexed with AMP-PNP (PDBID:4S32; 22) (Fig. S3), corroborating this conformational shift.

**Fig. 2:**
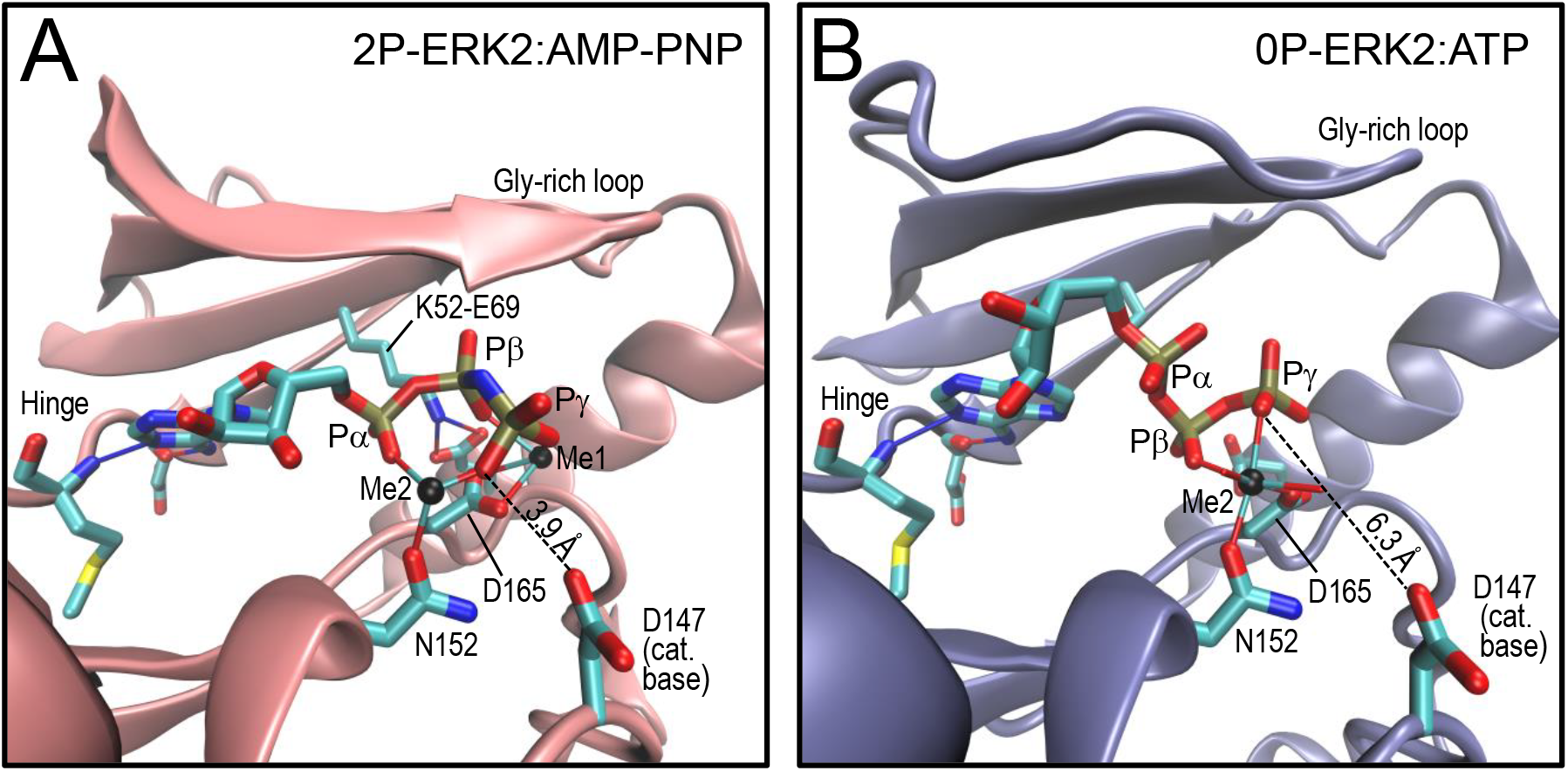
Crystal structures of ERK2 complexed with nucleotides. X-ray structures of **(A)** Mg^2+^-AMP-PNP complexed with 2P-ERK2 (2.9 Å), and **(B)** Mg^2+^-ATP complexed with 0P-ERK2 (PDBID:4GT3, ref. 20). The ribose ring shows distinctly different positioning in the two structures, resulting in a longer distance between Pγ-oxygens and the catalytic base in 0P-ERK2 compared to 2P-ERK2. Conserved motifs involved in nucleotide binding include the catalytic base (D147), Mg^2+^-coordinating residues (N152 and D165 in the DFG motif), K52-E69 salt bridge, the hinge region and the Gly-rich loop.

Detailed comparison of the structures in Fig. 2 showed closure between N- and C-terminal domains, forming closer interactions with nucleotide in the 2P-ERK2 complex. Domain closure could be quantified by the dihedral angle between helices αC and αE (23), which differed by 5 degrees between the two complexes (Table S3), as well as by movement of the N-terminal Gly-rich loop towards the C-terminal domain (Fig. 2). In addition, conserved Lys52, which is disordered in the 0P-ERK2:ATP complex, is well-ordered and coordinated with Glu69 and Pα and Pβ oxygens in the 2P-ERK2:AMP-PNP complex (Fig. 2, Table S3). Thus, nucleotide-bound active ERK2 revealed closer domain and ion pair interactions within the catalytic site. Overall, the comparison between these structures suggests that the conformational exchange induced by activation of ERK2 allows a transition within the active site, from a non-productive nucleotide binding interaction with 0P-ERK2 to a productive binding mode in 2P-ERK2. We propose that the shift to the R state in 2P-ERK2 enables formation of a catalytically competent nucleotide complex.

### X-ray structures of ERK2 complexed with inhibitors

Next we determined X-ray structures of 2P-ERK2 complexed with high-affinity inhibitors. Dual phosphorylated ERK2 complexed with Vertex-11e (PDBID:6OPK) or GDC-0994 (PDBID:6OPH) underwent significant remodeling of the activation loop and L16 loop regions, mirroring conformational changes seen in the active apoenzyme (Figs. S1, S4). Both complexes were comparable to 2P-ERK2:AMP-PNP with respect to domain closure and the bidentate Lys52-Glu69 salt bridge (Fig. 3A,B, Table S3). Thus, the active site residues in the Vertex-11e and GDC-0994 co-crystals were aligned similarly to those in the productive kinase-nucleotide complex.

**Fig. 3:**
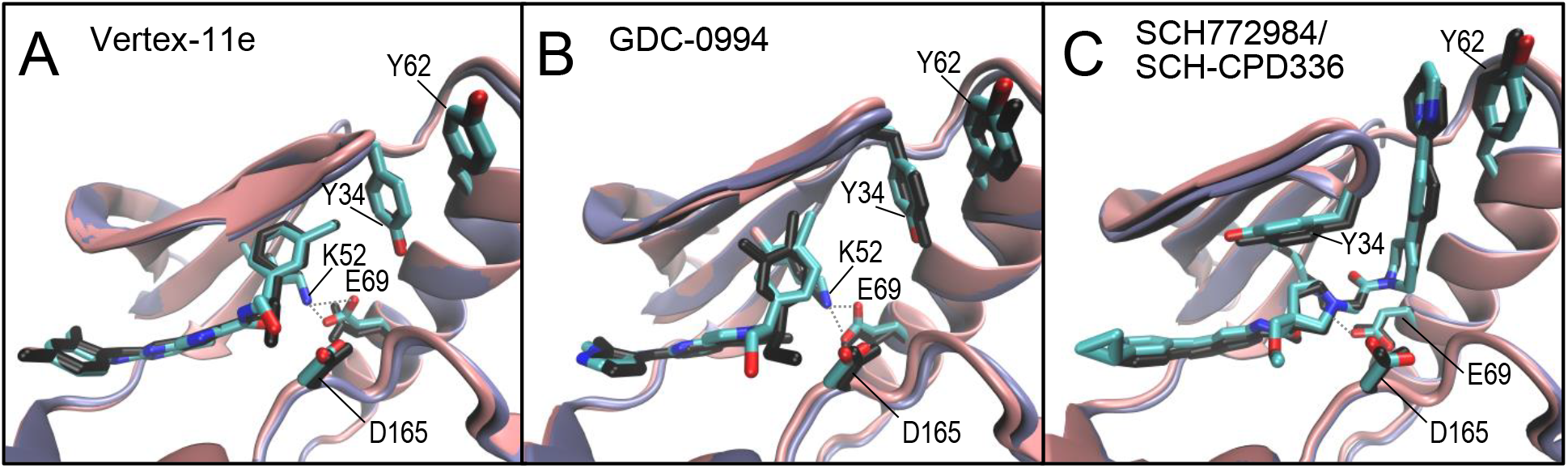
Crystal structures of ERK2 complexed with high affinity inhibitors. X-ray structures of the catalytic site in 2P-ERK2 (pink) and 0P-ERK2 (blue) complexed with **(A)** Vertex-11e or **(B)** GDC-0994. **(C)** Structures of 2P-ERK2 (pink) complexed with SCH-CPD336 and 0P-ERK2 (blue) complexed with SCH772984. The inhibitors adopt similar conformations within the catalytic sites of 2P-ERK2 (inhibitors colored by atom) and 0P-ERK2 (inhibitors colored black). Views of the full-length kinases are shown in Fig. S4.

SCH772984 is larger than Vertex-11e or GDC-0994, due to an extended piperazine-phenyl-pyrimidine motif (Fig. S2). In 0P-ERK2, this tricyclic motif disrupts stacking interactions between Y34 in the Gly-rich loop and Y62 in helix αC and forms new π-π stacking interactions between Y62 and the ligand (24). We solved an X-ray structure of 2P-ERK2 complexed with a close analogue of SCH772984, SCH-CPD336, which contains a partially saturated pyridine ring in place of the piperazine and two decorations at the opposite end of the molecule (Fig S2, PDBID:6OPI). SCH-CPD336 binds 2P-ERK2 with high affinity (K_d_ = 2.2 nM, measured by surface plasmon resonance, data not shown) and serves as a model for interactions between SCH772984 and the active kinase. As in the 0P-ERK2:SCH772984 complex, SCH-CPD336 disrupted Y34-Y62 stacking in 2P-ERK2, resulting in a monodentate Lys52-Glu69 interaction due to inhibitor-induced displacement of Lys52 (Fig. 3C, Table S3). The αC-αE dihedral angle in 2P-ERK2:SCH-CPD336 indicated an open conformation between N- and C-terminal domains, similar to that of 0P-ERK2:SCH772984, and closer to the apo- and nucleotide-bound forms of 0P-ERK2 than 2P-ERK2 (Table S3). Thus, 2P-ERK2 complexed with the SCH772984 analogue reveals a more open conformation compared to complexes with Vertex-11e and GDC-0994.

We compared these structures against reported inhibitor complexes with 0P-ERK2 (17,24). Nearly identical interactions were observed with both inactive and active forms of ERK2, with respect to inhibitor conformation and contacts with active site residues (Fig. 3, Fig. S4). Thus, unlike the nucleotide-bound complexes, structural differences between high affinity ligands bound to active *vs* inactive states of ERK2 were not detectable from X-ray data. Conceivably, the crystal forms may have masked differential binding modes through conformational trapping. Alternatively, the complexes might differ in conformational mobility around similar average conformations. Therefore, we hypothesized that solution measurements might be needed to observe differences corresponding to L and R states observed by NMR.

### HX-MS reveals differential inhibitor binding to the active site, coupled to the activation loop

HX-MS is a sensitive reporter of localized changes in structure and conformational mobility (25). We used HX-MS to examine effects of each inhibitor on deuterium uptake from solvent D_2_O, comparing active and inactive forms of ERK2 (Dataset S1). Inhibitor binding protected conserved regions in the active site pocket including the Gly-rich loop, helix αC, the hinge region, and the DFG motif (Fig. S5, S6), all of which were readily explained by proximity to the binding site. Long range effects were also indicated by protection of the αL16 helix (Fig. S5, S6). Deuterium uptake at the activation loop could not be compared directly between 0P and 2P kinase forms, due to differential proteolytic cleavage. However, structural changes could be monitored indirectly from the behavior of adjacent regions contacting the activation loop, including the P+1 substrate positioning loop and the N-terminal region of helix αF (Fig. S7).

The inhibitors altered deuterium uptake in two key regions that were instructive for understanding L and R conformations. The first was the DFG motif, which coordinates Mg^2+^ in the catalytic site. Here, AMP-PNP binding protected peptide LKICDFGL from hydrogen exchange to a greater degree in active ERK2 compared to inactive enzyme (Fig. 4A). The enhanced protection in 2P-ERK2 matched the X-ray evidence for a shift from nonproductive to productive nucleotide binding modes upon kinase activation, with greater steric protection from solvent due to closer interactions between Pγ/Mg^2+^ and DFG (cf. Fig. 2). GDC-0994 binding had comparable effects, also showing greater HX protection of the DFG region in active enzyme (Fig. 4B). Strikingly, Vertex-11e binding resulted in a significantly higher degree of protection, strongly interfering with deuterium uptake in 2P-ERK2 (Fig. 4C). Thus, the extent of HX protection of the DFG region was correlated with the degree of conformational selection for the R state in 2P-ERK2 complexed with AMP-PNP, GDC-0994 and Vertex-11e. By contrast, SCH772984, which is closer to the DFG motif due to its larger size, showed maximal HX protection in both 0P- and 2P-ERK2 (Fig. 4D). Overall, the varying patterns of protection of the DFG motif provided evidence for differential binding interactions between 0P-ERK2 and 2P-ERK2 for ligands able to form the R state in 2P-ERK2, whereas protection by SCH772984 was consistent with a single binding interaction.

**Fig. 4:**
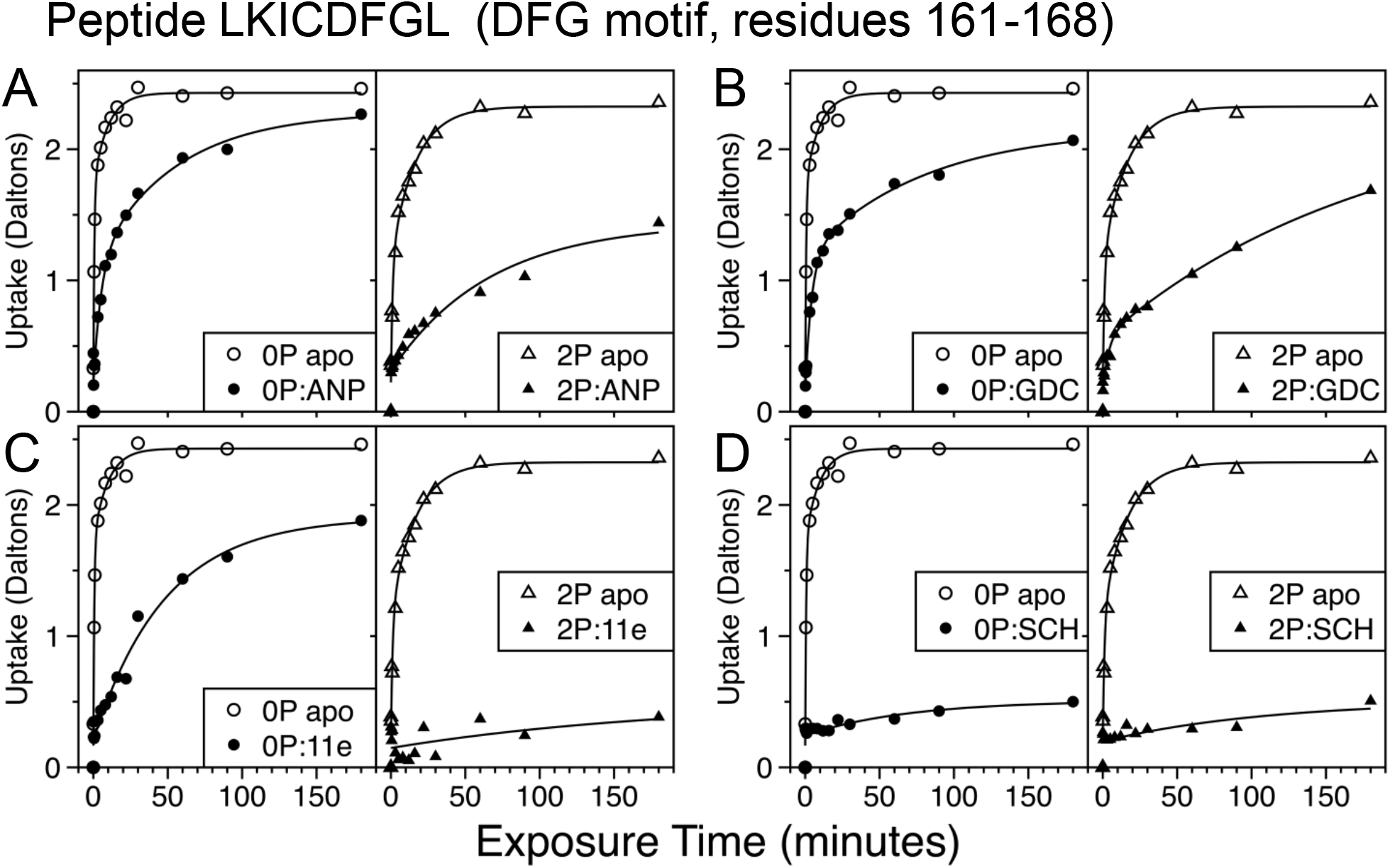
Inhibitors adopt distinct binding modes in the active sites of 2P- and 0P-ERK2. Closed symbols show HX-MS measurements of deuterium uptake into the DFG motif peptide, LKICDFGL, in 0P-ERK2 (circles) and 2P-ERK2 (triangles) complexed with **(A)** AMP-PNP, **(B)** GDC-0994, **(C)** Vertex-11e or **(D)** SCH772984. Open symbols show deuterium uptake into the DFG motif in 0P- or 2P-ERK2 apoenzyme forms. Binding of AMP-PNP, GDC-0994 or Vertex-11e leads to stronger protection of the DFG motif from solvent in active ERK2 than inactive kinase, correlating with degree of formation of the R state.

Deuterium uptake also reflected L and R states in the P+1 loop and αF N-terminus. This region is closely linked to the activation loop, and neither it nor the activation loop directly contact ligand (cf. Fig. S7). Deuterium uptake into peptides containing P+1 (YRAPEIML) and αF (YTKSIDI) showed greater protection in 2P-ERK2 than 0P-ERK2 apoenzyme (Figs. 5A, S8A), consistent with changes expected from structural remodeling of the activation loop. Likewise, with AMP-PNP or GDC-0994 bound, greater protection was observed in 2P-ERK2 than in 0P-ERK2, consistent with activation loop remodeling in both complexes (Figs. 5A,B, S8A,B). Importantly, Vertex-11e binding to 2P-ERK2 led to significantly greater HX protection of both peptides in the active kinase (Figs. 5C, S8C), while SCH772984 binding to 2P-ERK2 had the opposite effect, increasing deuterium uptake to a level higher than apoenzyme, and towards that seen with inhibitor-complexed 0P-ERK2 (Figs. 5D, S8D). Similar results were observed in a linker between helices αF and αG (peptide LSNRPIFPGKHYL) which forms hydrophobic and hydrogen bond interactions with the P+1 region (Figs. S7, S8E-H). To summarize, the HX protection of this key region is enhanced by phosphorylation and is consistent with folding of the activation loop into an active conformation in the 2P-ERK2 apoenzyme. Thus, increased HX protection by Vertex-11e binding and decreased protection by SCH772984 are respectively consistent with stabilization or destabilization of the active conformation of the activation loop, in a manner correlated with their conformational selection for R or L states.

**Fig. 5:**
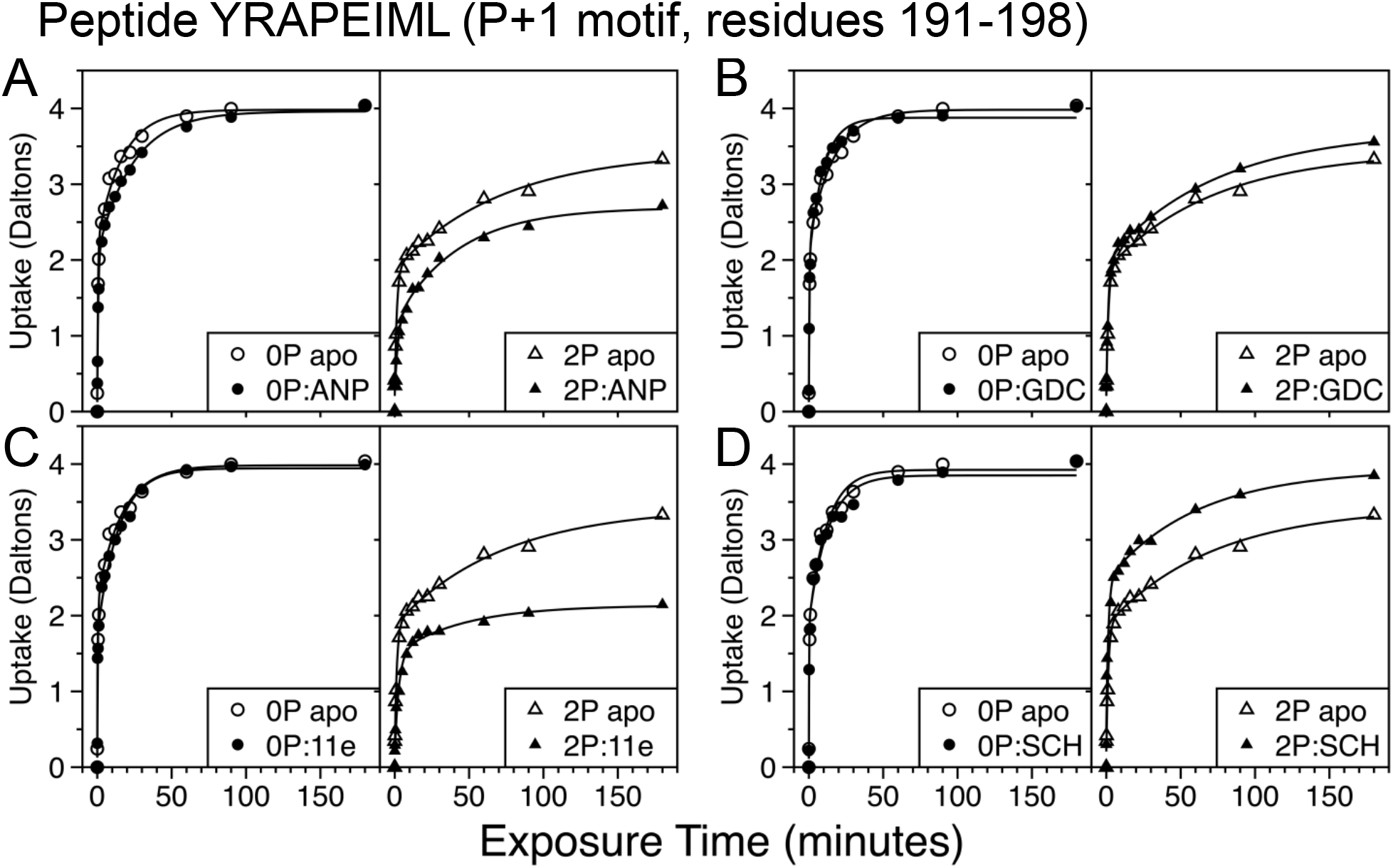
Vertex-11e and SCH772984 differentially regulate the activation loop in opposite directions. Effects of inhibitor binding on deuterium uptake into a region sensitive to the activation loop structure. Closed symbols show HX-MS measurements of deuterium uptake into the P+1 peptide, YRAPEIML, in 0P- and 2P-ERK2 complexed with **(A)** AMP-PNP, **(B)** GDC-0994, **(C)** Vertex-11e or **(D)** SCH774982. Open symbols show deuterium uptake into the P+1 motif in 0P- or 2P-ERK2 apoenzymes. The phosphorylation-induced HX protection of the P+1 motif in apo 2P-ERK2 reports folding of the activation loop into an active conformation. The increased HX protection by Vertex-11e binding and decreased protection by SCH772984 are consistent with stabilization or destabilization, respectively, of the active conformation.

### Inhibitor binding modulates ERK dephosphorylation

The HX results revealed that Vertex-11e and SCH772984 control conformational changes at the activation loop, suggesting that the activation loop is coupled to global exchange within the kinase core. Specifically, the results imply that the dually phosphorylated activation loop undergoes motions that track the L⇌R equilibrium. We considered the possibility that conformation-selective inhibitors alter the dynamics of the activation loop in a manner that affects regulatory mechanisms for ERK2. Previous studies have reported effects of Vertex-11e and SCH772984 on the steady state phosphorylation of ERK1/2 in human cancer cells (17,24,26). Often, SCH772984 treatment suppresses dual phosphorylation of ERK1/2, while Vertex-11e increases phosphorylation, without affecting the activity state of MKK1/2. Therefore, we investigated whether these inhibitors could affect dephosphorylation of 2P-ERK2 catalyzed by the MAP kinase phosphatase, MKP3/DUSP6. As demonstrated previously (27) and confirmed by MS/MS sequencing (data not shown), MKP3 dephosphorylates pY185 first to yield pT183-monophosphorylated (1P) ERK2, followed by removal of pT183 to yield 0P-ERK2. Therefore, time courses were fit to apparent first order rate constants for stepwise dephosphorylation of 2P- to 1P-ERK2 (k_1_) and 1P- to 0P-ERK2 (k_2_). Vertex-11e binding decreased k_1_ 2.5-fold compared to apoenzyme, while SCH772984 binding increased k_1_ 3-fold (Fig. 6, Table S4). Neither inhibitor significantly affected k_2_. GDC-0994 does not alter phosphorylation of cellular ERK1/2 (26), and neither it nor AMP-PNP affected MKP3-catalyzed dephosphorylation of 2P-ERK2 (Table S4). The outcomes are consistent with a model where conformational selection for the R state by Vertex-11e prevents motions of the activation loop by trapping it in the active conformation, thus blocking access of pY185 to MKP3. Conversely, conformational selection for the L state by SCH772984 increases accessibility of the activation loop to dephosphorylation. Thus, by regulating motions at the activation loop, inhibitors with properties of conformational selection can impact the recognition of ERK2 by MAP kinase phosphatase.

**Fig. 6:**
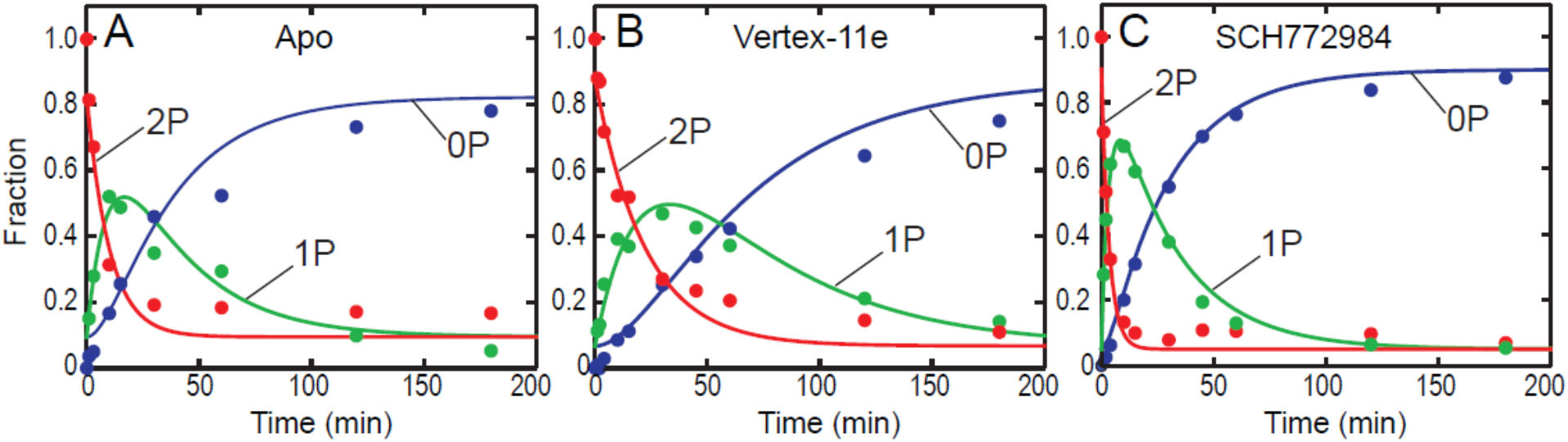
Vertex-11e and SCH772984 alter MKP3-catalyzed dephosphorylation of 2P-ERK2. MKP3 was incubated with 2P-ERK2 for indicated times, and mass spectrometry was used to quantify 2P, 1P (monophosphorylated-pT183) and 0P forms, in **(A)** apoenzyme, **(B)** Vertex-11e bound, and **(C)** SCH772984-bound ERK2. First order rate constants and errors for stepwise dephosphorylation events are presented in Table S4. Vertex-11e binding decreases the rate constant for the conversion of 2P-ERK2 → 1P-ERK2 by two-fold, while SCH772984 increases it by three-fold.

## DISCUSSION

Our study demonstrates that tight-binding inhibitors of ERK2 exhibit conformational selection for distinct allosteric states that are involved in L⇌R interconversion. By combining solution measurements and X-ray crystallography to monitor conformational changes induced by these inhibitors, we show that L and R states correspond to small but significant movements within the active site, which are coupled to larger motions of the activation loop. Our findings reveal that the activation loop undergoes exchange in the apo state of dual phosphorylated ERK2 and that motions of the activation loop can be controlled allosterically by inhibitor binding. As a consequence, inhibitors that bind with conformational selection can control the accessibility of the pT183 and pY185 phosphorylation sites to MAP kinase phosphatase. Thus, we propose that the dynamics of the activation loop in 2P-ERK2 function to facilitate its recognition by regulatory proteins, such as MAP kinase phosphatases (Fig. 7).

**Fig. 7:**
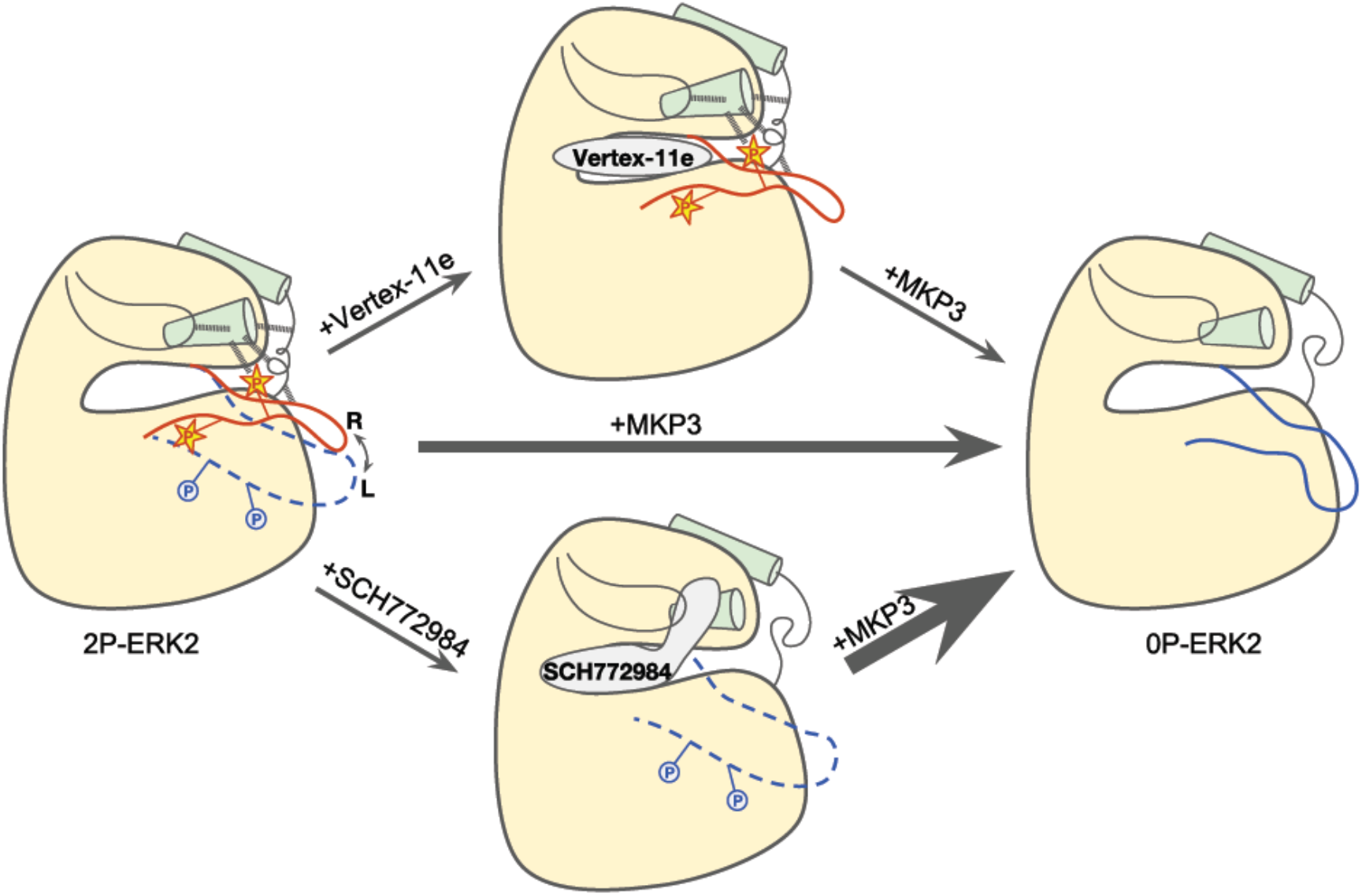
Model for allosteric control of activation loop dynamics by conformation-selective inhibitors. The activation loop in 2P-ERK2 is dynamic, coupled to equilibrium exchange between L and R states, reflecting Gly-rich loop mobility and domain closure. Vertex-11e and SCH772984 exploit the low energetic barrier between L and R, to confer conformational selection for R and L, respectively. Their corresponding effects on activation loop motions slow or enhance dephosphorylation by MKP3.

Our findings provide insights into the conformational differences between L and R states. The NMR measurements of global exchange behavior at Ile, Leu, and Val methyls throughout the kinase core predict that the L and R states involve residues throughout the N- and C-domains and catalytic pocket. But while X-ray structures of 0P-ERK2 and 2P-ERK2 apoenzymes showed large changes at the activation loop and the L16 loop (Fig. S1), structural differences within the active site were less apparent. Our new structure of active, phosphorylated ERK2 complexed with AMP-PNP helps clarify these differences by revealing a nucleotide binding mode distinct from that of inactive, unphosphorylated ERK2 complexed with ATP. Here, the position of the Gly-rich loop in 0P-ERK2 allows a nucleotide conformation that places the Pγ-oxygen atoms >6 Å from the catalytic base. In the nucleotide complex with 2P-ERK2, closure of the Gly loop repositions the Pγ-oxygen closer to the catalytic base and also allows coordination of an additional Mg^2+^ ion, which is needed for full activity (28). Compared to 0P-ERK2:ATP, the 2P-ERK2:AMP-PNP complex also shows displacement of helix αC relative to helix αE, reflecting N-domain rotation and closure (23). These differences provide a structural interpretation of corresponding HX-MS measurements which reflect distinct binding interactions between AMP-PNP and the conserved DFG motif. It is significant that none of these changes alter the orientation of the DFG motif, which remains in the DFG-in state. Instead, the allosteric switch from L to R following ERK2 phosphorylation involves small differences in conformation of enzyme and nucleotide, leading to a shift from a nonproductive to a productive nucleotide binding mode. In this way, motions that allow L → R conversion can be associated with a shift to a catalytically competent state.

In contrast to AMP-PNP/ATP, high-affinity inhibitor co-crystal structures displayed nearly identical interactions within the active sites of 2P- and 0P-ERK2. All structures displayed the DFG-in state, unlike what is observed in structures of kinases complexed with conventional Type II inhibitors (16). Nevertheless, measurements by HX-MS clearly showed that, like AMP-PNP, Vertex-11e and GDC-0994 have different binding modes depending on kinase activity state. Their ability to form the R state in 2P-ERK2, measured by NMR, correlated well with their degree of DFG protection, measured by HX-MS. These results suggest that the ability of 2P-ERK2 to access the R state enables differential binding to DFG by AMP-PNP and GDC-0994, as well as conformational selection by Vertex-11e. Thus, solution measurements reported conformational shifts within the active site that were masked in the crystal forms. These shifts describe an exchange equilibrium that appears distinct from the DFG-in/DFG-out switch, but can nevertheless confer Type II-like properties of conformational selection on ERK inhibitors.

Outside the catalytic site, ERK2 activation induced larger conformational changes. Vertex-11e or GDC-0994 binding to 2P-ERK2 allowed remodeling of the activation loop and L16 region, similar to apoenzyme. In contrast, SCH-CPD336 bound to 2P-ERK2 showed disorder in the activation loop and L16 regions, likely due to the disruption of Gly-rich loop and αC interactions by the piperazine-phenyl-pyrimidine moiety. These structural changes agreed well with differences in hydrogen exchange protection in the P+1 and αF N-terminal region, which reports conformational change at the activation loop. Importantly, the enhanced degree of HX protection by Vertex-11e compared to AMP-PNP or GDC-0994 matched that expected for its shift of the L⇌R equilibrium further towards the R state. These findings suggest coupling between inhibitor occupancy of the active site and the distally-located activation loop.

Taken together, the solution measurements and crystal structures are consistent with a model in which dynamics of the activation loop are associated with the equilibrium between L and R in 2P-ERK2. This model differs from structural analyses suggesting an activation loop conformation in the active kinase which is stabilized by an extensive salt bridge network between pT183, pY185 and five Arg sidechains (R68, R146, R170, R189, R192) (5). Conceivably, favorable enthalpic contributions from these salt bridges could be counteracted by a loss in entropy due to residue immobilization (29). This may explain the small difference in free energy (~0.8 kcal/mol) favoring R over L, measured by NMR (11). The low energetic barrier allows Vertex-11e to trap the active conformation of the activation loop, and SCH772984 to disrupt it.

The ability of conformation-selective inhibitors to control motions of the phosphorylated activation loop is an emerging concept for the kinase field. Recent studies have documented activation loop motions in different protein kinases. For example, altered motions due to Type II inhibitor binding have been shown by site-directed spin labeling EPR measurements of the activation loop in p38α MAPK (30). Likewise, activation loop dynamics have been suggested by simulations of the Fyn tyrosine kinase domain (31). However to date, only Aurora A kinase has been found to undergo activation loop motions in its phosphorylated form (32–35). Here, time-resolved FRET and single-molecule fluorescence spectroscopic measurements show interconversion of the phosphorylated activation loop between active and inactive conformations. These are coupled to exchange between DFG-in and DFG-out conformations, and can be shifted by Type II inhibitors (34,35). The characteristics of ERK2 differ from Aurora A, in that dynamic motions of the activation loop coupled to global exchange in the active site involve movements distinct from DFG backbone rotation.

Our investigations provide evidence for the ability of ERK inhibitors to control activation loop motions and in doing so, regulate kinase recognition by phosphatases. This represents new evidence that conformation-selective inhibitors can allosterically control the regulation of wild type ERK. Interestingly, mutant forms of ERK1/2 containing amino acid substitutions at the gatekeeper residue, Q103A/T, and a residue adjacent to the DFG motif, C1164L, allow binding by conventional Type II inhibitors (35,36). Here, crystal structures of the unphosphorylated ERK2 double mutant revealed that Type II inhibitors bound to a DFG-out conformer. They also affected the activation loop conformation and blocked dephosphorylation by MKP3. These inhibitors were unable to bind non-mutant ERK2, providing evidence that the kinase in its wild-type form cannot access the DFG-out state. The findings indicate that restraints in ERK2 that prevent DFG backbone rotation can be found in specific side chain packing interactions within the regulatory spine. We propose that motions in the activation loop can be coupled to different types of conformational changes in the active site, ranging from large backbone rotations to form DFG-out conformers, induced by mutation of unphosphorylated ERK2, to small changes involving differential ligand interactions with the DFG-in state, which can be induced by phosphorylation of wild-type ERK. How the architecture of ERK2 controls its dynamics and allosteric regulation is a fascinating question for future exploration.

## MATERIALS AND METHODS

Vertex-11e, SCH772984, and GDC-0994 were obtained from Selleck Chemicals and adenylyl-imidodiphosphate (AMP-PNP) was from Sigma-Aldrich. SCH-CPD336 was synthesized in-house, following methods described in US Patent WO2007070298A1, example 336. 0P-ERK2 and 2P-ERK2 [*methyl*-^1^H,^13^C]-ILV labelled at ILV for NMR or unlabelled for HX-MS were prepared as described (11,14,15). Detailed methods for measurements by NMR, HX-MS and X-ray crystallography followed previously described protocols. They are described in detail in Suppl. Information, which includes Suppl. Materials and Methods, Figures and Datasets.

## ACKNOWLEDGEMENTS

We are indebted to Drs. Thomas Lee and Danijel Djukovic for assistance with LC-MS instrumentation and analysis, to Dr. Marcelo Sousa and Sandra Metzner, for the gift of His_6_-SUMO-ratERK2, and to the 2015 CCP4/APS summer school program for use of the Argonne National Laboratory beamline and help with X-ray data collection. This work was supported by NIH grant awards R01GM114594 and S10RR026641 (NGA), and T32GM008759 (JCL and DBI). JCL gratefully acknowledges support by the Eugene Huffman Memorial Scholarship and the Sheryl R. Young Memorial Scholarship, Univ. Colorado.

