## Supplemental Methods, Tables, and Figures for "Activation loop dynamics are controlled by conformation-selective inhibitors of ERK2"

### PEGRAM ET AL, SUPPLEMENTAL INFORMATION

#### SUPPLEMENTAL MATERIALS AND METHODS

#### SUPPLEMENTAL REFERENCES

#### SUPPLEMENTAL TABLES

Table S1. Kinetic parameters for ERK2 inhibitors

Table S2. X-ray data collection and refinement parameters

Table S3. Dihedral angles between $\alpha$C-$\alpha$E and Lys52-Glu69 distances

Table S4. Rate constants for dephosphorylation of 2P-ERK2 by MKP3

#### SUPPLEMENTAL FIGURES

Fig. S1. Crystal structures of ERK2 apoenzymes.

Fig. S2. Chemical structures of ligands used in this study.

Fig. S3. Corroborating crystal structures of nucleotide-bound ERK2.

Fig. S4. Crystal structures of ERK2 complexed with high affinity inhibitors.

Fig. S5. Proteolytic peptides analyzed by HX-MS.

Fig. S6. Deuterium uptake into regions of ERK2 altered by ligand binding.

Fig. S7. Peptide reporters of activation loop conformation.

Fig. S8. Deuterium uptake plots for peptide reporters of activation loop conformation.

#### SUPPLEMENTAL DATASETS

Dataset S1. Deuterium uptake time courses for all HX-MS experiments (Excel format).

#### SUPPLEMENTAL MATERIALS AND METHODS

**Materials.** Vertex-11e, SCH772984, and GDC-0994 were obtained from Selleck Chemicals. SCH-CPD336 was synthesized in-house, following methods described in US Patent WO2007070298A1, example 336. Adenylyl-imidodiphosphate (AMP-PNP), dimethylsulfoxide (DMSO), dithiothreitol (DTT), and formic acid were from Sigma-Aldrich; and $D_2O$ (99.9%) was from Cambridge Isotope Laboratories. Ni-NTA agarose was obtained from Qiagen.

**Protein Preparations.** NMR experiments used wild-type $His_6$-ratERK2 selectively labeled with [methyl-$^1H$,$^{13}C$]Ile, Leu, and Val (ILV) as described (1). HX-MS and X-ray crystallography complexes with Vertex-11e used wild-type rat ERK2 with an N-terminal $His_6$-SUMO tag and C-terminal ULP1-cleavable linker expressed from a pET32 plasmid, constructed by Marcelo Sousa and Sandra Metzner, U. Colorado, Boulder. $His_6$-SUMO-ERK2 was induced in *E. coli* BL21(DE3) cells with 0.5 mM isopropyl β-D-1-thiogalactopyranoside (IPTG) for 19 h at 18°C. Cells were resuspended in lysis buffer (50 mM potassium phosphate pH 8.0, 300 mM NaCl) with cOmplete EDTA-free protease inhibitor cocktail (Roche), lysed by sonication, and centrifuged at 18,000 rpm for 1 h. Supernatant was filtered (0.45 μm) and loaded onto a gravity-flow column with 3 mL Ni-NTA resin equilibrated in lysis buffer + 10 mM imidazole, and protein was eluted with lysis buffer + 0.5 M imidazole. EDTA (10 μM) and ULP1 protease (~3 μg) were added, followed by overnight dialysis into lysis buffer at 4°C. Cleaved protein was then reloaded onto a gravity-flow column with 3 mL Ni-NTA resin and recovered with 2-4 column volumes of lysis buffer + 10 mM imidazole. Unphosphorylated ERK2 (0P-ERK2) was further purified by anion exchange FPLC (1 mL HiTrap Q HP, GE Healthcare), eluting in two peaks using a linear gradient from 50-300 mM NaCl in 50 mM Tris, pH 7.4, 1 mM DTT. Only the first peak (eluting at 130 mM NaCl) was used for analysis (2). Half of the purified protein was used as 0P-ERK2, and the other half was dual-phosphorylated to 2P-ERK2 with constitutively active MKK1-G7B and purified by anion exchange FPLC as described (3). Purified 0P- and 2P-ERK2 were dialyzed into HX buffer and stored in aliquots at -70°C. Phosphorylation stoichiometries of 0P-ERK2 and 2P-ERK2 were respectively >95% 0P and >96% 2P, as determined by LC-MS/MS (4) using a Waters Synapt G2 mass spectrometer in positive ion mode with PLGS 3.0 software.

X-ray structures of 2P-ERK2 complexed with AMP-PNP, GDC-0994 and SCH-CPD336 used $His_6$-EK-humanERK2 induced in *E. coli* BL21(DE3) cells with 1 mM IPTG for 17 h at 20°C. Cells were resuspended in lysis buffer (20 mM Tris pH 8.0, 10%(v/v) glycerol, 10 mM imidazole, 7.5 mM 2-mercaptoethanol) with 300 mM NaCl and cOmplete EDTA-free protease inhibitor cocktail (Roche), lysed by mechanical disruption, and clarified by centrifugation at 30,000xg for 40 min. Supernatant was batch loaded onto Talon Metal Affinity Resin for 1 h at 4°C. Resin was washed with lysis buffer and subsequently eluted with lysis buffer + 100 mM imidazole. Talon elution was pooled and diluted with QAE buffer A (20 mM Tris pH 8.5, 10% glycerol, 7.5 mM 2-mercaptoethanol) and loaded onto a Source 15Q anion exchange column (GE Healthcare). The column was developed using a linear gradient of 0-350 mM NaCl in QAE buffer, collecting only the first peak (eluting at 175 mM) for further purification. 0P-ERK2 protein was diluted to 1 mg/mL, cleaved using EKMax Enterokinase (Invitrogen), and dual-phosphorylated to 2P-ERK2 with constitutively active MKK1 (MEK1ca-6His(62-393). 2P-ERK2 was batch purified with Ni-NTA and EKAway resins (Invitrogen) to remove unclipped ERK2, MKK1 and EKMax. The unbound fraction containing clipped 2P-ERK2 was diluted with QAE buffer A and purified by Source 15Q anion exchange chromatography as described for 0P-ERK2. Pooled 2P-ERK2 was concentrated and further purified by Superdex 200 SEC equilibrated with 20 mM HEPES pH 7.5, 150 mM NaCl and 10 mm DTT. 2P-ERK2 was concentrated and stored in aliquots at -70°C.

MAP kinase phosphatase-3 (MKP3/DUSP6) was expressed and purified as described (5). BL21(DE3)-pLysS cells transformed with plasmid pMCSG7-MKP3 (Addgene#40772, ref 6) were grown in 2x YT medium at 37°C and induced with 0.2 mM IPTG for 4 h at 22°C. Cells were disrupted in lysis buffer (50 mM Tris, pH 7.4, 300 mM NaCl, 10 mM 2-mercaptoethanol, EDTA-free protease inhibitor cocktail) by sonication. Supernatants were clarified by centrifugation (18,000 rpm, 1 h) and filtration (0.45 μm), incubated by batch with 6 mL Ni-NTA resin (1 h), washed with 50 mM Tris pH 6.0, 400 mM NaCl, 10 mM imidazole, 10 mM 2-mercaptoethanol, 10%(v/v) glycerol, and eluted with the same buffer containing 50, 100, 300 and 500 mM imidazole. MKP3 was pooled from 100 and 300 mM imidazole fractions and concentrated with buffer exchange into 25 mM Tris pH 8.0, 250 mM NaCl,10 mM 2-mercaptoethanol, 1 mM EDTA. Enzyme was quantified by absorbance at 280 nm and stored at -70°C.

**Enzyme Assays.** Measurements of kinetic parameters ($K_d$, $k_{on}$, $k_{off}$) for GDC-0994 binding to 2P-ERK2 were performed as previously described (7). Dephosphorylation assays were performed with 4 mM 2P-ERK2 and 200 nM MKP3 in buffer containing 100 mM sodium acetate, 50 nM Tris, 50 mM Bis-Tris, pH 6.5, incubated at 25 °C for up to 3 h. For each time point, 50 mL aliquots were removed and the reaction terminated by adding formic acid to 1% (v/v). Potassium hydroxide was added to adjust the pH to 7.4, 0.5 μg modified trypsin (Promega) was added, and samples were proteolyzed at 37 °C overnight. 2P-, 1P- and 0P-ERK2 were quantified by LC-MS as described (4), using a Waters Synapt G2 mass spectrometer with PLGS 3.0 software to monitor $MH_2^{+2}$ and $MH_3^{+3}$ ions corresponding to peptide VADPDHDHTGFLTEYVATR. First order rate constants were obtained by fitting the resulting data to a model for sequential dephosphorylation (2P→1P→0P) using Kintek Explorer (Kintek Corp.).

**NMR Spectroscopy.** [*methyl*-$^1$H,$^{13}$C]Ile, Leu, and Val-labeled 0P- and 2P-ERK2 were prepared as described (1) in buffer containing 50 mM Tris pH 7.4, 150 mM NaCl, 5 mM $MgSO_4$, 0.1 mM EDTA, 5 mM DTT, 100%(v/v) $D_2O$, and 2.5% (v/v) glycerol. SCH772984 and GDC-0994 stock solutions (2.5 mM) were prepared in DMSO-d6, due to limited solubility in $D_2O$, and added to 2P-ERK2 (40 μM) or 0P-ERK2 (30 μM), to form complexes with 100% binding stoichiometry ([inh]: [protein] = 1.2:1). The DMSO-d6 concentration in the final protein sample was ~1%(v/v).

Two-dimensional (2D) $^{13}$C–$^1$H heteronuclear multiple quantum coherence (HMQC) spectra of 0P- and 2P-ERK2 were collected on Varian VNMRS 800 and 900 MHz NMR spectrometers. Data were collected at 25°C for a total time of 8 h for 0P-ERK2 and 11 h for 2P-ERK2. Each spectrum was acquired with 144 (800 MHz) or 160 complex points (900 MHz) in the $t_1$ ($^{13}$C) dimension, corresponding to 28.8 ms at 800 MHz and 29.1 ms at 900 MHz, and 1024 complex points in the acquisition period. WURST40 $^{13}$C decoupling was applied during the 85 ms acquisition period, and a 1.5 s delay period was used between each scan. The spectral processing was performed with the software package NMRPipe (8). Time domain data in the $^1$H dimension were apodized by a cosine-squared window function and zero-filled prior to Fourier transformation. The indirect dimension ($^{13}$C) was apodized by a cosine window function and zero-filled prior to Fourier transformation. Spectral visualization and analysis were achieved using CCPNMR Analysis software (9).

**X-ray Crystallography.** All X-ray structure images were created with VMD software (10). Cocrystals of 2P-ERK2:Vertex-11e mixed 2P-ERK2 [300 nmol in 50 mL of column buffer (25 mM HEPES pH 7, 150 mM NaCl, 1 mM TCEP)] with equimolar Vertex-11e (300 nmol in 0.5 mL DMSO), adding inhibitor in aliquots of 50 μL while stirring. The protein-inhibitor complex was concentrated by binding onto a 1 mL Mono-Q column and step-eluting with 2 mL column buffer + 0.5 M NaCl. The complex was then purified by size-exclusion chromatography (S-200 in column buffer) and concentrated to 10 mg/mL protein. The complex was crystallized from hanging drops at room temperature, mixing 1 µL protein + 1 µL well solution (16.5% PEG3350, 100 mM Tris-HCl pH 8.5). Crystals were soaked in cryoprotectant (20% PEG3350, 100 mM Tris-HCl pH 8.5) prior to freezing and transport. Data was collected at the Advanced Photon Source at Argonne National Laboratory (Lemont, IL). The raw electron diffraction data were processed with DIALS (11). Refinement and model building were performed using RefMac, Phenix, and Coot.

Co-crystals of 2P-ERK2 in complex with AMP-PNP, GDC-0994 or SCH-CPD336 were each grown by vapor diffusion. Complexes were set up by mixing 10 mg/mL 2P-ERK2 with 0.8 mM $Mg^{2+}$-AMP-PNP, 1 mM GDC-0994, or 1 mM SCH-CPD336, and incubated for 60 min at room temperature. Complexes were then clarified by centrifugation and supernatants recovered. Crystals of 2P-ERK2:AMP-PNP were grown at 20°C from sitting drops in 24 wells, mixing 1.0 µL protein + 1.0 µL well solution (36% PEG3350, 0.1 M Tris pH 8.8). One round of macroseeding was required to grow crystals large enough for data collection. Crystals of 2P-ERK2:SCH-CPD33 were grown at 20°C from sitting drops in 96 wells, mixing 0.5 µL protein + 0.5 µL well solution (20% (w/v) PEG3350, 100 mM Na cacodylate pH 7.2, 600 mM NaCl, 10 mM $MnCl_2$). Crystals of 2P-ERK2:GDC-0994 were crystallized from hanging drops at 20°C by mixing 0.9 µL protein + 0.9 µL well solution (36% PEG3350, 100mM bicine pH 9.0). Crystals of 2P-ERK2:AMP-PNP and 2P-ERK2:SCH-CPD336 were cryoprotected in well solution + 20%(v/v) ethylene glycol, and crystals pf 2P-ERK2:GDC-0994 were cryoprotected in well solution + 10% glycerol prior to data collection. X-ray diffraction data were collected in-house on a Rigaku FR-E generator with Osmic confocal mirrors and an Raxis-IV++ detector. The structure was solved with Phaser, Refmac and Coot (CCP4 suite) (12).

**Hydrogen Exchange Mass Spectrometry (HX-MS).** 2P- or 0P-ERK2 (10-12 μM) was incubated with 15 μM inhibitor or 4 mM AMP-PNP for 30 min in HX buffer (50 mM potassium phosphate pH 7.2, 100 mM NaCl, 5 mM DTT, 10 mM $MgCl_2$ and 0.5% (v/v) DMSO from concentrated ligand solutions). Deuterium uptake was initiated by adding 90 μL $D_2O$ (containing an equal concentration of ligand, DMSO, and $MgCl_2$) to 10 μL ERK2, to reach a final buffer concentration of 5 mM potassium phosphate pH 7.2, 10 mM NaCl, 0.5 mM DTT, 10 mM $MgCl_2$, and 0.5% DMSO. Reactions were incubated at 25.0°C for varying times between 30 s and 180 min. Exchange was quenched with 80 μL 100 mM potassium phosphate pH 2.2, and 100 μL was immediately injected onto a Waters nanoAcquity HDX Manager UPLC system for proteolysis on an immobilized pepsin column (Poroszyme, Applied Biosystems) at 10°C. Proteolysis, as well as peptide desalting (0°C, Vanguard Acquity UPLC BEH C18), were carried out in isocratic Solvent A (0.1%(v/v) formic acid in $H_2O$) with flowrate 100 μL/min. Peptide separations (Waters Acquity UPLC BEH C18, 1.7 μm, 1.0x100 mm) were carried out using a 12 min linear gradient from 8-85% Solvent B (0.1% formic acid in acetonitrile) at 40 μL/min.

Peptides were analyzed by ESI-MS/MS using a Waters Synapt G2 HDMS Q-TOF mass spectrometer in positive ion mode. Continuum data were collected over 50-2000 m/z with 0.23 s scan time. Lock mass correction was achieved using Glu-Fibrinogen peptide ($MH_2^{+2}$ = 785.8426 m/z), with resolution = 20,000 at m/z 956. Undeuterated pepsin-cleaved peptides from 0P- or 2P-ERK2 were identified by $MS^e$ sequencing using PLGS 3.0 (Waters). Searches of rat ERK2 allowed nonspecific digestion and variable modifications of oxidized Met and phosphorylated Ser/Thr/Tyr, with parameter settings of intensity threshold = 750 counts, lock mass window = 0.4 Da, and low energy threshold = 135 counts. Peptides were accepted after filtering for high confidence, which required maximum $MH^+$ error 15 ppm, minimum sequence length 3, maximum sequence length 25, minimum product ions 3, and identification in at least 2 of 3 replicate runs. Deuterium uptake was calculated for high confidence peptides in each hydrogen exchange dataset using DynamX 3.0 (Waters). Raw files were processed using a 0.35 Da lock mass window and a low energy threshold of 100. For every experiment, automated ion assignments for deuterated peptides were manually inspected and validated. The weighted average mass of isotope distributions was calculated for each verified peptide using DynamX, and referenced to the weighted average mass of its unlabeled form.

HX reproducibility was evaluated by measuring four independent time courses of 2P-ERK2 apoenzyme, which were performed over 14 months by two separate operators using two separate enzyme preparations. Measured for 10 time points and 92 peptides, the average standard deviation of deuterium uptake was 0.10 Da (S.E.M. = 0.05 Da) with average coefficient of variation = 0.029.

#### Table S1. Kinetic parameters for ERK2 inhibitors

| | 0P-ERK2 | 2P-ERK2 | | |
|---|---|---|---|---|
| Inhibitor | $K_d$ (nM) ± s.d. | $k_{on}$ ($M^{-1}s^{-1}$) ± s.d. | $k_{off}$ ($s^{-1}$) ± s.d. | $K_d$ (nM) ± s.d. |
| Vertex-11e [a] | 2.5 ± 0.5 | $0.21 \times 10^{-6}$ ± $0.04 \times 10^{-6}$ | $5.8 \times 10^{-5}$ ± $1.9 \times 10^{-5}$ | 0.21 ± 0.04 |
| GDC-0994 | n.d. | $1.9 \times 10^{-6}$ ± $0.1 \times 10^{-6}$ | $330 \times 10^{-5}$ ± $30 \times 10^{-5}$ | 1.7 ± 0.1 |
| SCH772984 [a] | 0.12 ± 0.04 | $0.12 \times 10^{-6}$ ± $0.04 \times 10^{-6}$ | $31 \times 10^{-5}$ ± $9 \times 10^{-5}$ | 1.2 ± 0.10 |

[a] Data from Rudolph et al. (2015) *Biochemistry* 54:22-31 (ref 13).

#### Table S2. X-ray data collection and refinement parameters

| CRYSTAL STRUCTURE | 2P:AMP-PNP | 2P:GDC-0994 | 2P:SCH-CPD336 | 2P:VERTEX-11E |
|---|---|---|---|---|
| **PDB CODE** | 6OPG | 6OPH | 6OPI | 6OPK |
| **DATA COLLECTION:** | | | | |
| SPACE GROUP | $P2_12_12_1$ | $P2_12_12_1$ | $P2_12_12_1$ | $P2_12_12_1$ |
| UNIT CELL PARAMETERS (Å,°) | a=41.9, b=77.2, c=152.2, β=90.0 | a=42.1, b=76.9, c=151.8, β=90.0 | a=41.4, b=77.6, c=126.5, β=90.0 | a=42.0, b=78.1, c=152.6, β=90.0 |
| RESOLUTION RANGE (Å) | 23.9-2.90 2.97-2.90 | 26.6-2.40 2.46-2.40 | 22.1-3.00 3.08-3.00 | 30.0-2.54 2.61-2.54 |
| UNIQUE REFLECTIONS | 11508 | 17814 | 8550 | 19482 |
| COMPLETENESS (%) | 99.37 | 88.67 | 99.67 | 99.90 |
| **REFINEMENT:** | | | | |
| $R_{WORK}/R_{FREE}$ (%) | 15.6/23.9 | 19.3/24.3 | 18.5/26.0 | 19.3/24.1 |
| NO. OF PROTEIN ATOMS | 2858 | 2854 | 2662 | 2839 |
| NO. OF WATERS | 159 | 17 | 30 | 20 |
| NO. OF HETEROATOMS | 33 | 31 | 50 | 32 |
| R.M.S.D. BONDS (Å) | 0.009 | 0.007 | 0.008 | 0.008 |
| R.M.S.D. ANGLES (°) | 1.7 | 1.6 | 1.7 | 1.7 |
| CRUICKSHANK DPI (Å) | 0.37 | 0.36 | 0.45 | 0.38 |
| **AVERAGE B FACTORS (Å$^2$):** | | | | |
| PROTEIN | 52.5 | 60.4 | 50.6 | 79.4 |
| WATERS | 43.6 | 43.6 | 33.6 | 71.4 |
| LIGAND | 48.4 | 60.5 | 39.1 | 65.2 |

#### Table S3. Dihedral angles between αC-αE and Lys52-Glu69 distances

| Crystal structures | Helix αC-αE dihedral angles [a] | Lys52-Glu69 distances (Å) (Lys-NZ-Glu-OE1; Lys-NZ-Glu-OE2) |
|---|---|---|
| 2P-ERK2 Apo (2ERK) | 95° | 3.4; 3.9 Å |
| 0P-ERK2 Apo (5UMO) | 88° | 4.5; 4.8 Å |
| 2P-ERK2:AMP-PNP (6OPG) | 94° | 2.9; 2.9 Å |
| 0P-ERK2:ATP (4GT3) | 90° | --- [b] |
| 2P-ERK2:AMP-PCP (5V60) | 95° | 3.1; 3.8 Å |
| 0P-ERK2:AMP-PNP (4S32) | 88° | 3.7; 5.1 Å |
| 2P-ERK2:Vertex-11e (6OPK) | 97° | 2.6; 3.2 Å |
| 0P-ERK2:Vertex-11e (4QTE) | 94° | 2.9; 3.2 Å |
| 2P-ERK2:GDC-0994 (6OPH) | 94° | 2.7; 3.1 Å |
| 0P-ERK2:GDC-0994 (5K4I) | 94° | 2.7; 3.0 Å |
| 2P-ERK2:SCH-CPD336 (6OPJ) | 89° | 3.1; 5.0 Å |
| 0P-ERK2:SCH772984 (4QTA) | 90° | 2.7; 4.7 Å |

[a] The dihedral angle between helix αC (residues 62-75) and helix αE (residues 121-140) was calculated using HackaMol version 0.051 (13). The two best-fit vectors along the backbone atoms of each helix were used to generate the four points at +/- ½ the corresponding Cα to Cα distance from the geometric center of each helix. Increased angles correspond to domain closure between N- and C-terminal lobes.

[b] Not measurable due to disorder of Lys52 side chain.

#### Table S4. Rate constants for dephosphorylation of 2P-ERK2 by MKP3

Model: $\text{2P-ERK2} \xrightarrow{k_1} \text{1P-ERK2(pT183)} \xrightarrow{k_2} \text{0P-ERK2}$

| | $k_1$ ($min^{-1}$) ± S.E. | $k_2$ ($min^{-1}$) ± S.E. |
|---|---|---|
| Apo | 0.10 ± 0.04 | 0.03 ± 0.02 |
| Vertex-11e | 0.04 ± 0.01 * | 0.02 ± 0.01 |
| GDC-0994 | 0.13 ± 0.02 | 0.05 ± 0.02 |
| SCH772984 | 0.29 ± 0.03 ** | 0.03 ± 0.01 |
| AMP-PNP | 0.07 ± 0.01 | 0.10 ± 0.03 |

Rate constants in the presence of inhibitors were not significantly different from Apo ($p > 0.05$), except for $k_1$, Vertex-11e (*$p = 0.011$) and $k_1$, SCH772984 (**$p = 0.0006$).

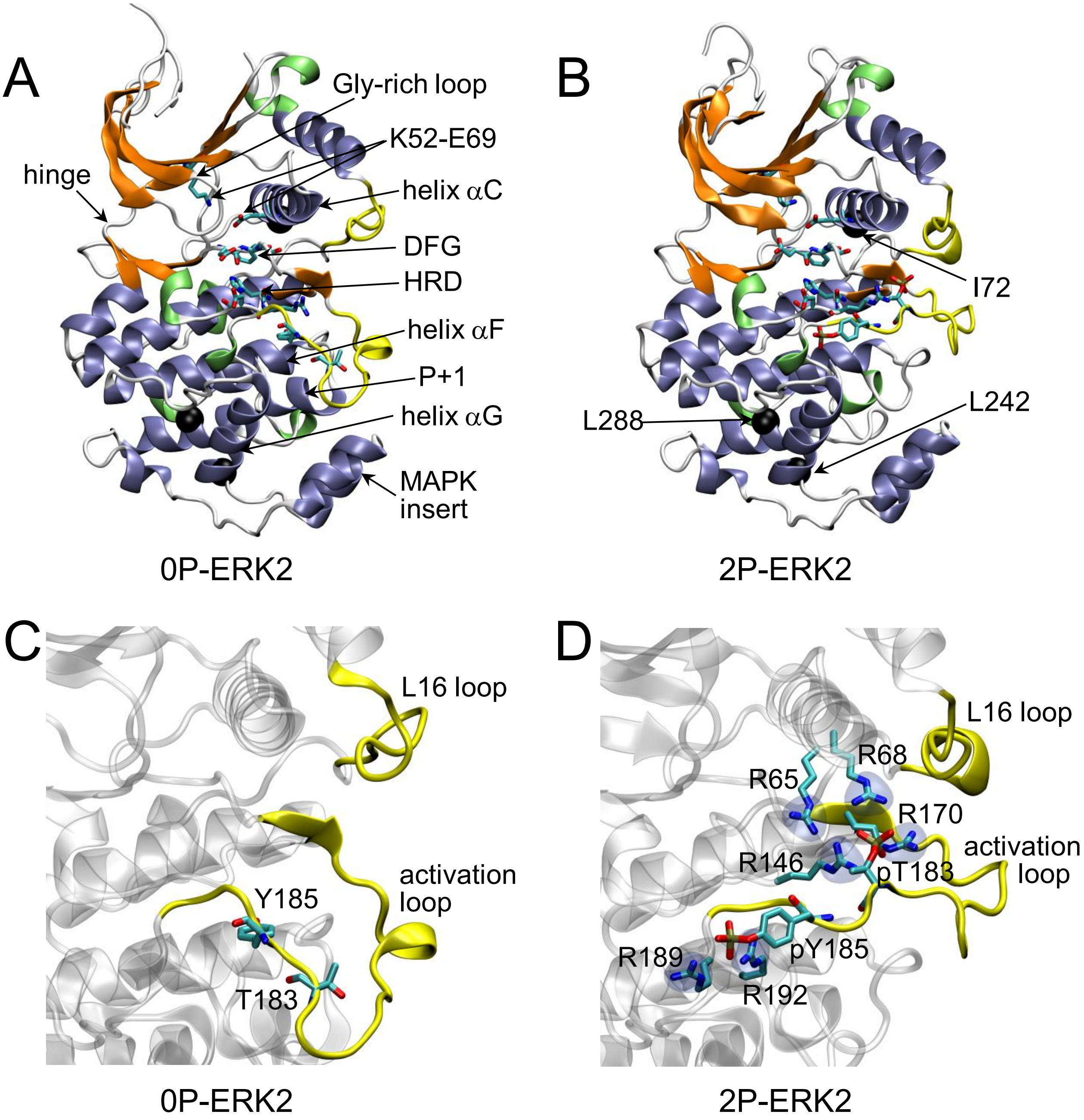

**Fig. S1. Crystal structures of ERK2 apoenzymes. (A,B)** X-ray structures of **(A)** 0P-ERK2 (PDBID:5UMO) and **(B)** 2P-ERK2 (PDBID:2ERK). Panel **A** labels conserved motifs in ERK2 common to protein kinases. Panel **B** labels residues which illustrate L⇌R exchange in HMQC spectra shown in Fig. 1. **(C,D)** The largest conformational changes accompanying dual phosphorylation at T183 and Y185 occur at the activation loop and L16 loop regions, shown in **(C)** 0P-ERK2 and **(D)** 2P-ERK2. In 2P-ERK2, pT183 and pY185 form ion pairs with multiple Arg residues, while the L16 loop folds into a 3/10 helix and forms new interactions with the activation loop.

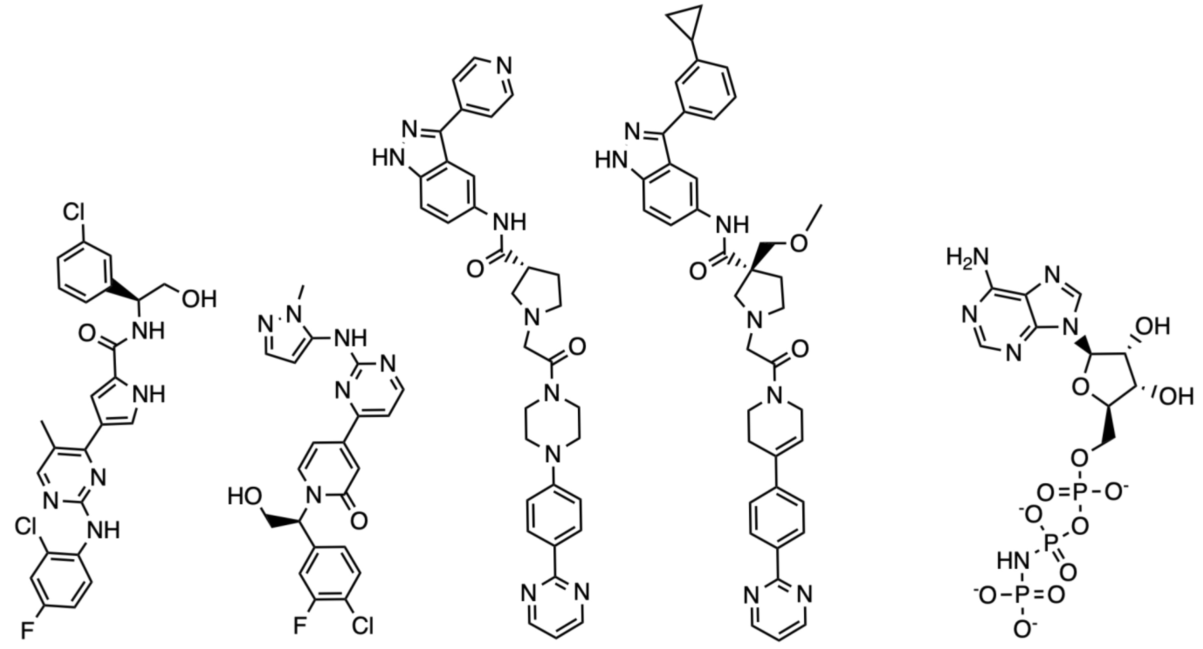

A. Vertex-11e B. GDC-0994 C. SCH772984 D. SCH-CPD336 E. AMP-PNP

**Fig. S2. Chemical structures of ligands used in this study. (A)** Vertex-11e, **(B)** GDC-0994, **(C)** SCH772984, **(D)** SCH-CPD336 (analogue of SCH772984) and **(E)** AMP-PNP.

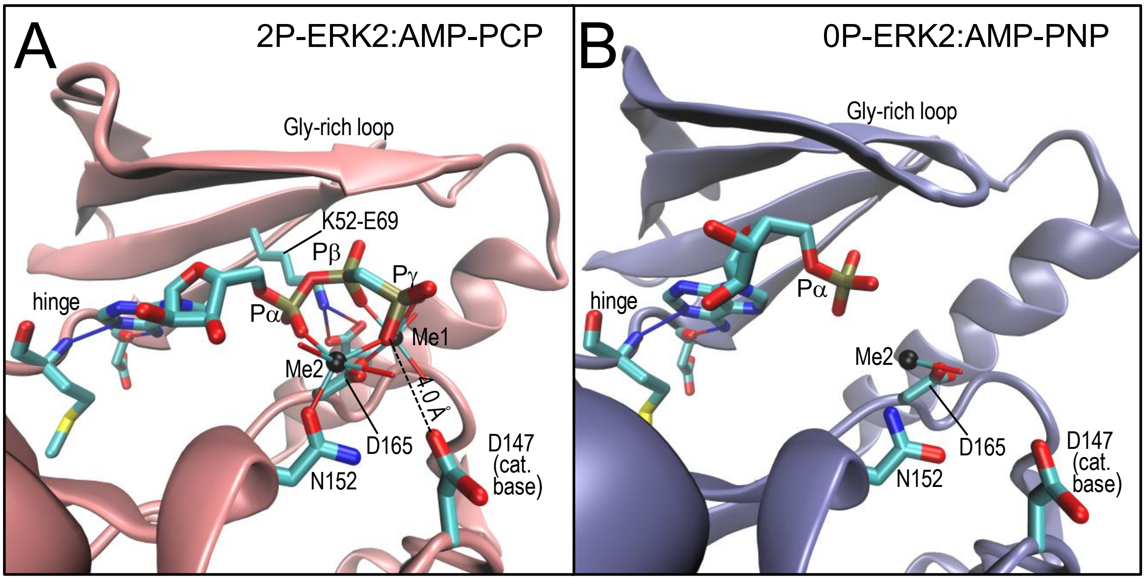

**Fig. S3. Corroborating crystal structures of nucleotide-bound ERK2.** X-ray structures of **(A)** 2P-ERK2 complexed with AMP-PCP (PDBID:5V60, ref 21) and **(B)** 0P-ERK2 complexed with AMP-PNP (PDBID:4S32, ref 22). Although incomplete due to disorder, these structures corroborate the conformational shifts in the ribose dihedral angle between 2P-ERK2 and 0P-ERK2 observed in Fig. 2.

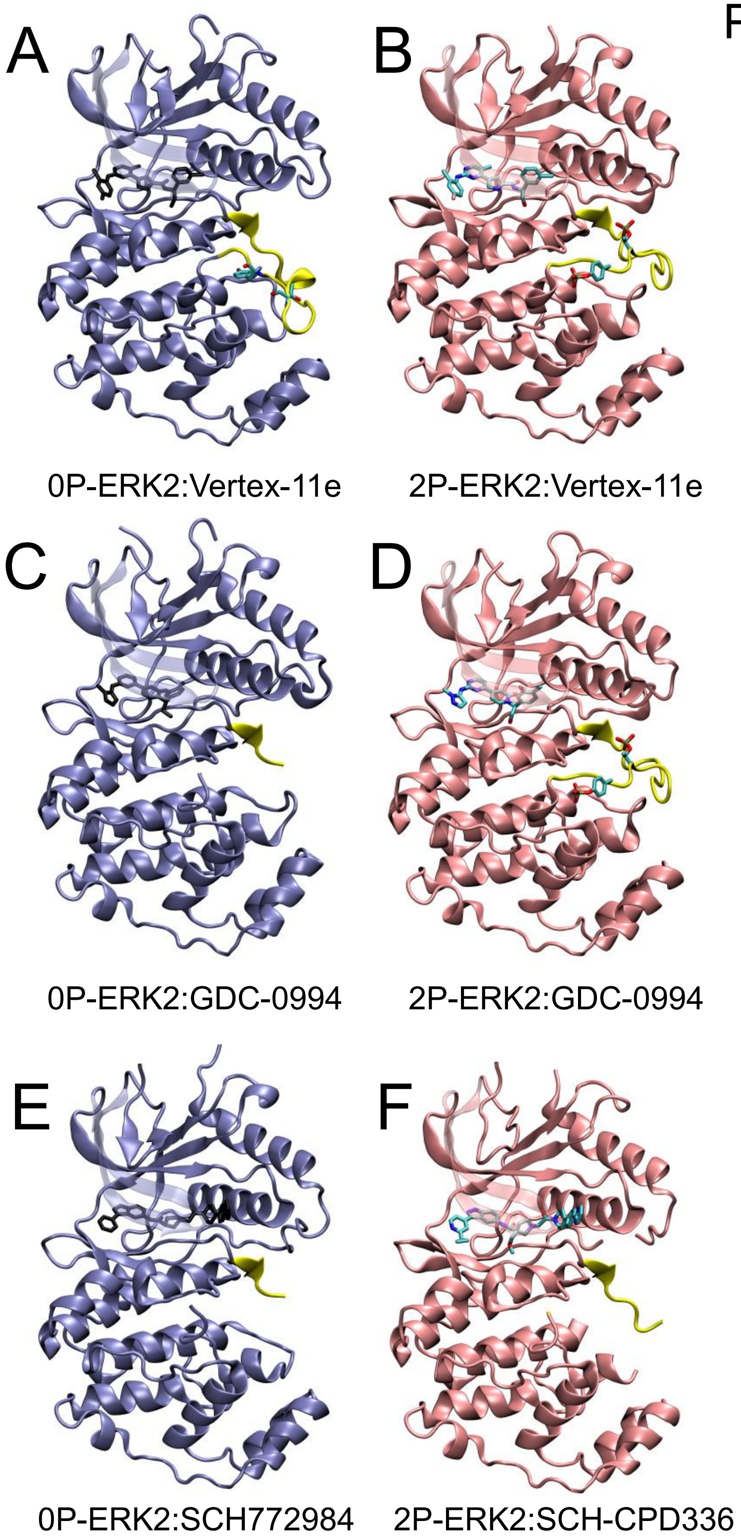

**Fig. S4. Crystal structures of ERK2 complexed with high affinity inhibitors.** Full view of ERK2 X-ray structures shown in Fig. 3. **(A,B)** 0P- and 2P-ERK2 complexed with Vertex-11e, **(C,D)** 0P- and 2P-ERK2 complexed with GDC-0994, **(E)** 0P-ERK2 complexed with SCH772984, and **(F)** 2P-ERK2 complexed with SCH-CPD336. Activation loop remodeling of 2P-ERK2 is comparable to apoenzyme when complexed with Vertex-11e or GDC-0994. Activation loop was not observed in 0P-ERK2 complexed with SCH772984 and 2P-ERK2 complexed with SCH-CPD336.

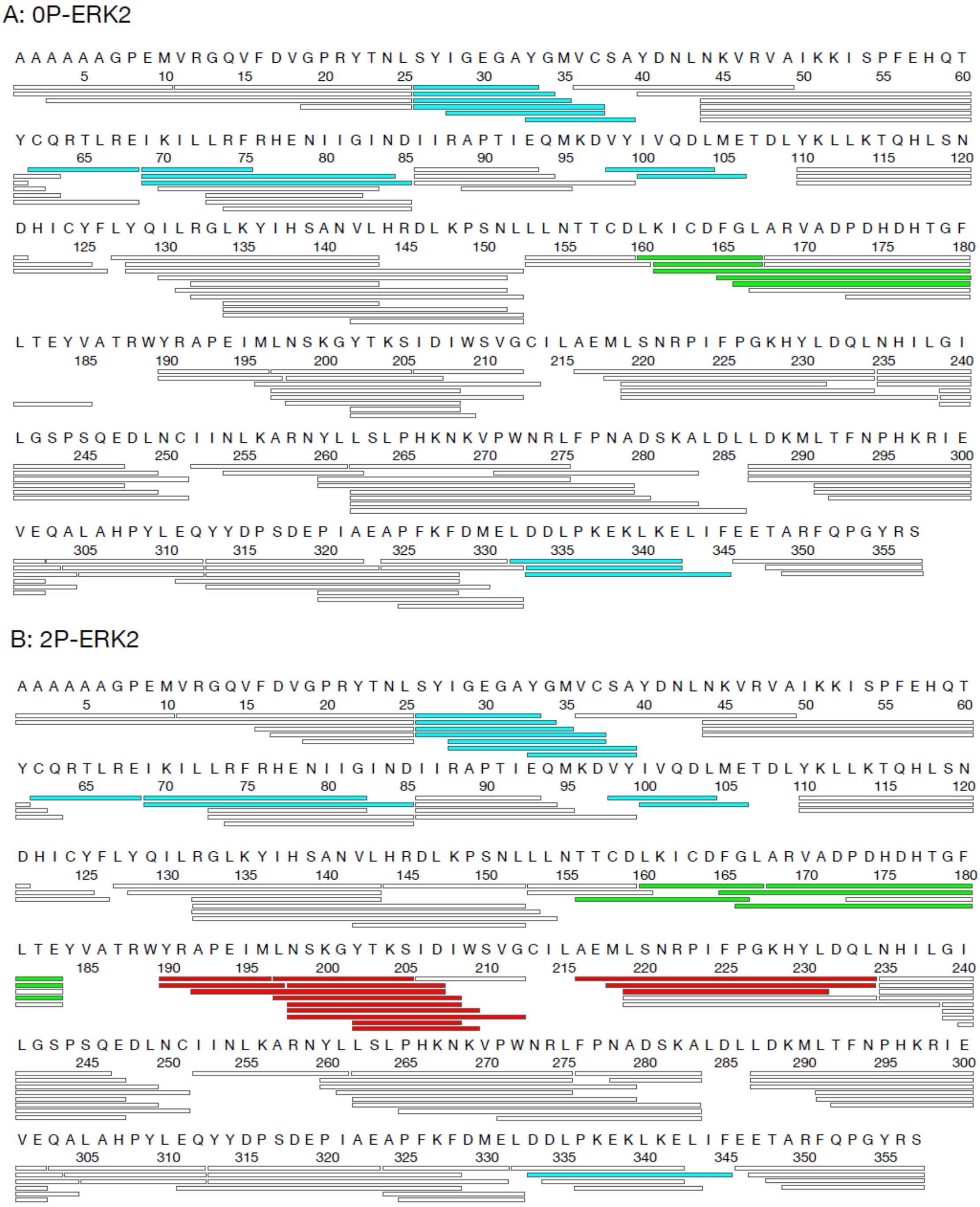

**Fig. S5. Proteolytic peptides analyzed by HX-MS.** Peptides produced by pepsin digestion yield 94% and 93% coverage of exchangeable amides in **(A)** 0P-ERK2 and **(B)** 2P-ERK2, respectively. Differential proteolysis precludes direct comparison of the activation loop between 0P- and 2P-ERK2. Colors indicate regions where ligand binding alters HX. Blue indicates peptides where ligand binding results in similar HX protection of 0P- and 2P-ERK2 (see Fig. S6). Green and red indicate peptides respectively at the DFG motif and near the activation loop, where ligands showed differential protection of 0P- and 2P-ERK2 (see Figs. 4, 5 and S8). Regions of HX protection or enhancement by ligand binding were identified by ≥4 consecutive time points with differences from apo of ≥0.4 Da. These peptides are shown in Fig S6. Other overlapping peptides that showed the same patterns are also indicated on the coverage map.

A: YGMVCSA (res. 34-40, Gly-rich loop)

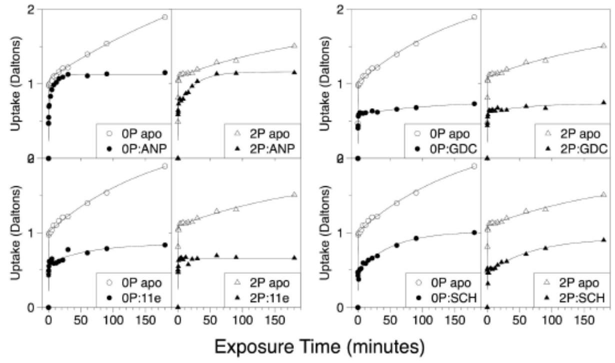

B: CQRTLRE (res. 63-69, Helix $\alpha$C)

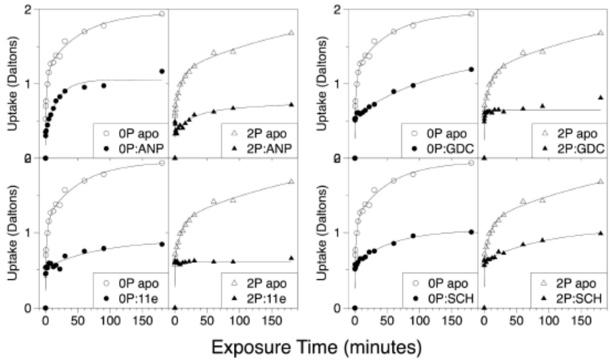

C: VYIVQDL (res. 99-105, $\beta$5–Hinge)

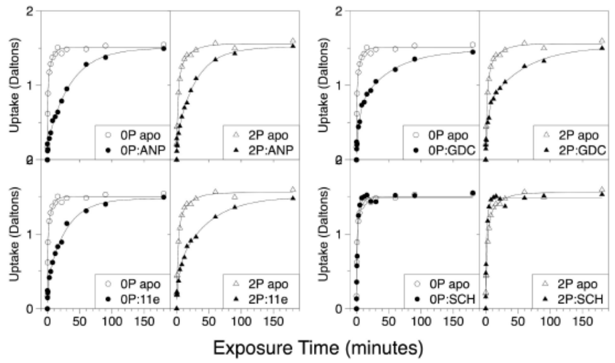

D: IVQDLME (res. 101-107, Hinge)

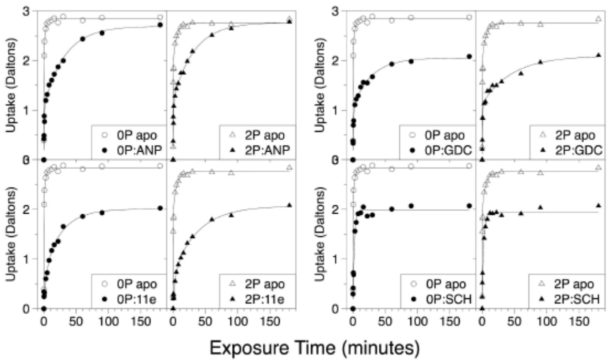

E: DDLPKEKLKELIF (res.334-346, helix $\alpha$L16)

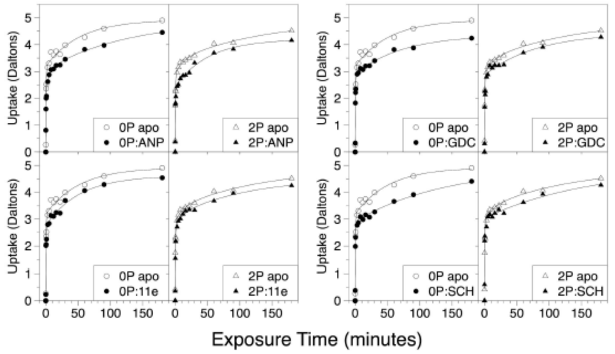

**Fig. S6. Deuterium uptake into regions of ERK2 altered by ligand binding.** Peptides representative of conserved kinase sequence motifs at or near the catalytic pocket of ERK2 include: **(A)** the Gly-rich loop (34-40), which contacts nucleotide phosphate oxygens; **(B)** helix $\alpha$C (63-69), containing E69 which forms a conserved salt bridge with $\beta$3-K52 to coordinate ATP; **(C,D)** the hinge region, which coordinates the adenine ring in ATP (99-105 and 101-107); **(E)** helix $\alpha$L16, which interacts with helix $\alpha$C (334-347). Each region shows HX protection in 0P-ERK2 (circles) and 2P-ERK2 (triangles), when complexed with AMP-PNP, GDC-0994, Vertex-11e, and SCH772984. Ligand bound forms are shown with closed symbols. Apoenzyme forms are shown with open symbols.

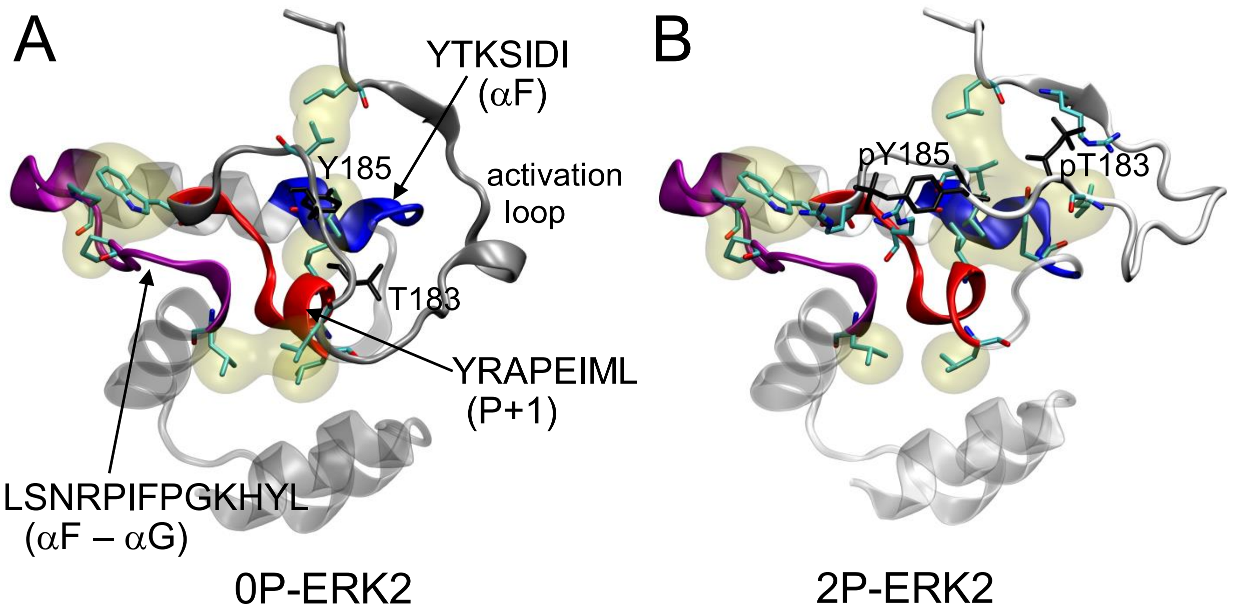

**Fig. S7. Peptide reporters of activation loop conformation.** Peptides from the P+1 loop (YRAPEIML) and the αF N-terminus (YTKSIDI) form direct contacts with the activation loop, while the αF-αG loop (LSNRPIFPGKHYL) forms hydrophobic and hydrogen bond interactions with the P+1 loop. Each region shows greater HX protection in 2P-ERK2, reflecting the activation loop remodeling between **(A)** 0P-ERK2 and **(B)** 2P-ERK2.

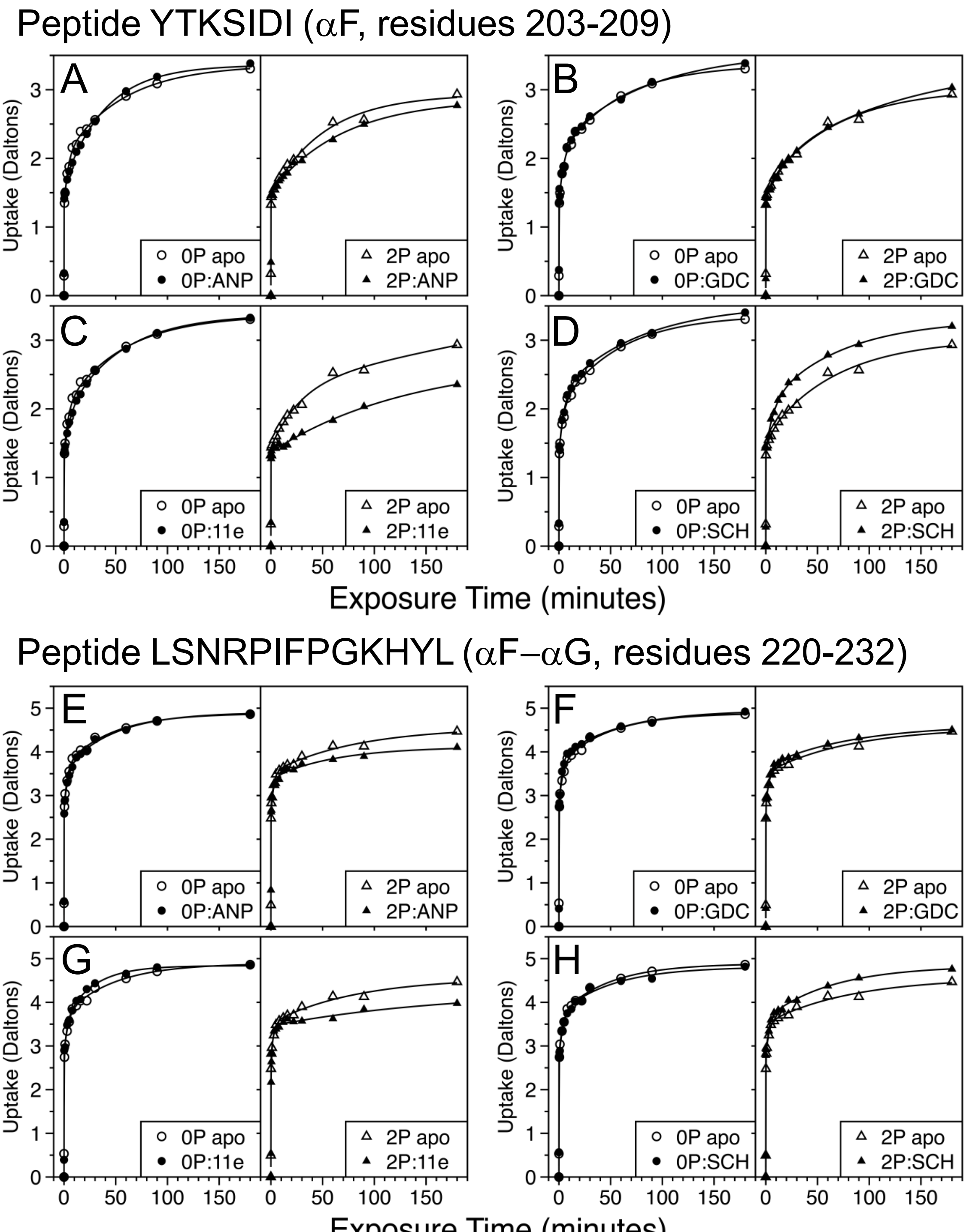

**Fig. S8. Deuterium uptake plots for peptide reporters of activation loop conformation.** Effects of inhibitor binding on deuterium uptake into **(A-D)** the αF N-terminal peptide (YTKSIDI), and **(E-H)** the αF-αG loop peptide (LSNRPIFPGKHYL). Both regions are sensitive to the activation loop structure, and corroborate the behavior of the P+1 peptide (YRAPEIML) shown in Fig. 5. Closed symbols show HX-MS measurements of deuterium uptake in 0P- and 2P-ERK2 complexed with **(A,E)** AMP-PNP (ANP), **(B,F)** GDC-0994, **(C,G)** Vertex-11e or **(D,H)** SCH772984. Open symbols show deuterium uptake in 0P- or 2P-ERK2 apoenzymes.
